## Supplementary Figures, Tables, and Notes for "FtsZ treadmilling is essential for Z-ring condensation and septal constriction initiation in *Bacillus subtilis* cell division"

+Equal author contribution

### SUPPLEMENTARY FIGURES

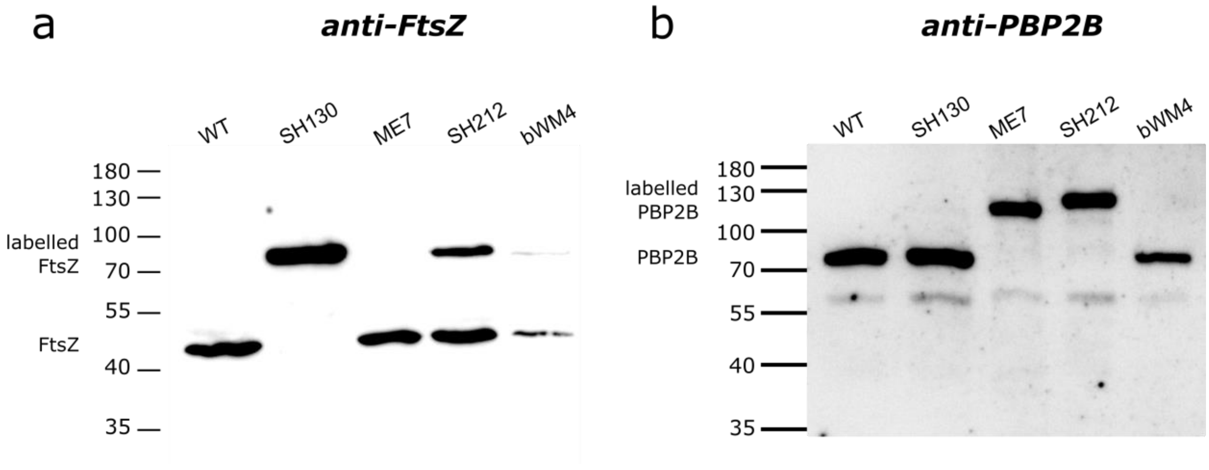


**Supplementary Figure 1: Western blots of fluorescent fusion proteins used in this study.** Strains with fusion proteins expressed from inducible promoters (SH212 and bWM4) were grown with the inducer concentration used in experiments (SH212: 20 µM IPTG, 0.08% xylose; bWM4: 25 µM IPTG). 20 µL lysate of each strain was blotted. Samples were incubated with specific polyclonal antibodies for FtsZ or PBP2B. Bands were visualised by a chemiluminescence system.

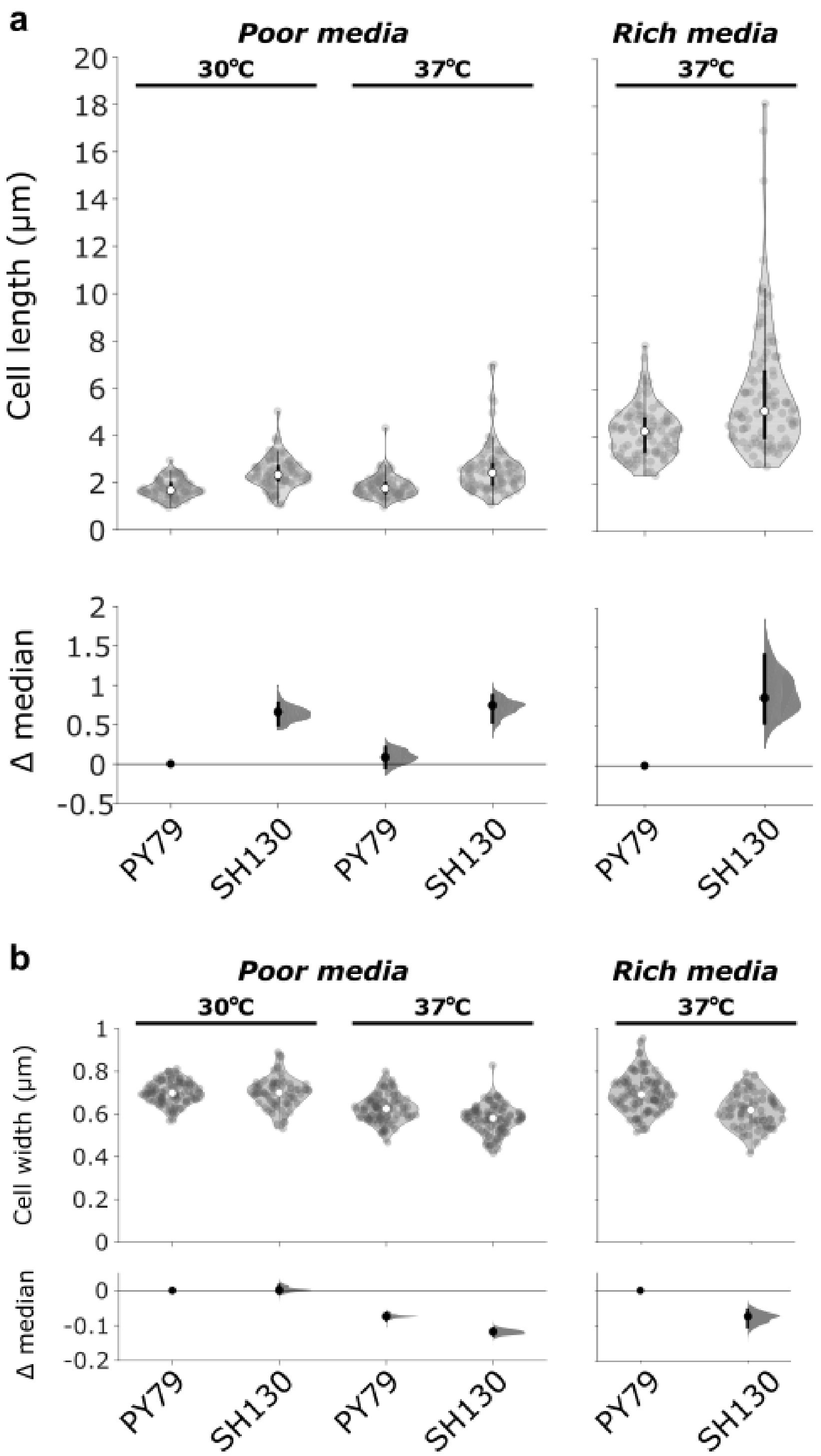


**Supplementary Figure 2: Cell length (a) and width (b) measurements for SH130 (expressing FtsZ-GFP at native locus) vs wild-type (PY79) in different growth conditions.** The cell length and width of SH130 was measured by microscopy of membrane stained cells in different growth conditions and compared to PY79. Cells were grown in poor (CDM) or rich media (PHMM) at the indicated temperature. Violin plots: white circles, median; thick black lines, interquartile range; thin black lines, 1.5x interquartile range. DABEST plots: black circle, median difference between indicated conditions; black lines, 95% confidence interval of median difference.

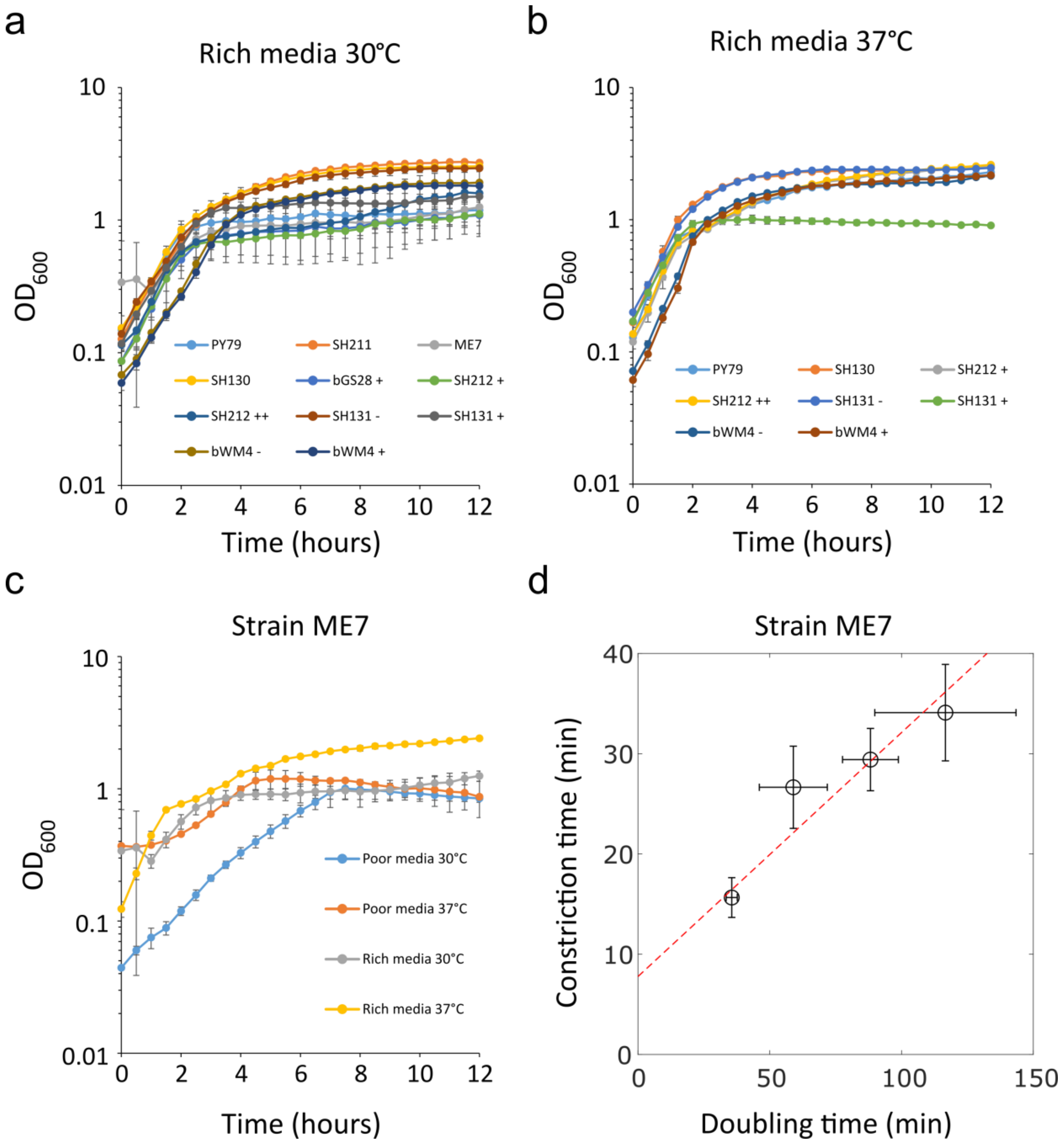


**Supplementary Figure 3: Growth curves of strains under different growth conditions.** Growth was monitored for 12 hours using a FluoStar plate reader (BMG Labtech) at the indicated temperature. Average values and standard deviations of triplicate repeats are plotted. (a-b) Mutant strains grown in rich media (PHMM) at low (a) and high (b) temperatures compared to wild-type (PY79). Strains containing inducible promoters are displayed both with (+) and without (-) inducer. Since strain SH212 has two inducible promoters, it is shown both as IPTG induction only (+) or IPTG and xylose induction (++). Inducer concentrations are as follows: bGS28, 20 µM IPTG; SH212, 20 µM IPTG and 0.08% xylose; SH131, 10 µM IPTG; bWM4, 25 µM IPTG. (c) Strain ME7 grown in both rich (PHMM) and poor (CDM) media at two different temperatures. (d) Relation between doubling times in liquid culture and measured constriction times for ME7 strain. Error bars are standard deviation. Red dotted line: linear fit to data.

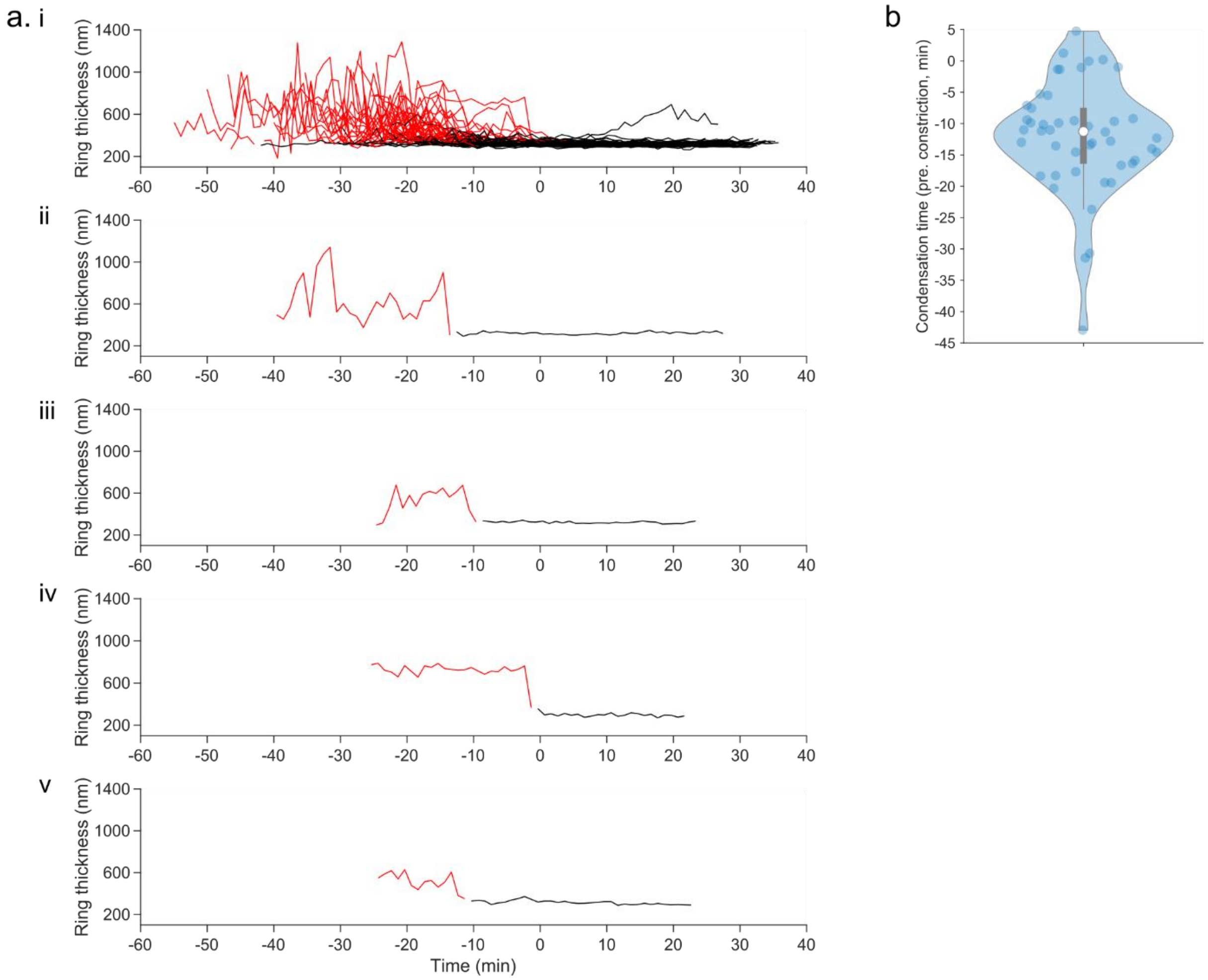


**Supplementary Figure 4: Change point analysis of FtsZ-ring condensation.** (a) All classified time series of Z-ring axial thickness for cells expressing FtsZ-GFP (SH130), corresponding to data shown in Figure 1b. (i) shows all time traces, (ii-v) shows 4 exemplar traces, same traces as highlighted in Figure 1b. Red line indicates decondensed state assigned by changepoint detection, black line indicates condensed state. (b) Violin plot of observed condensation time for all axial thickness time traces presented in (a) where 2 states were detected. White circles, median; thick grey lines, interquartile range; thin grey lines, 1.5x interquartile range.

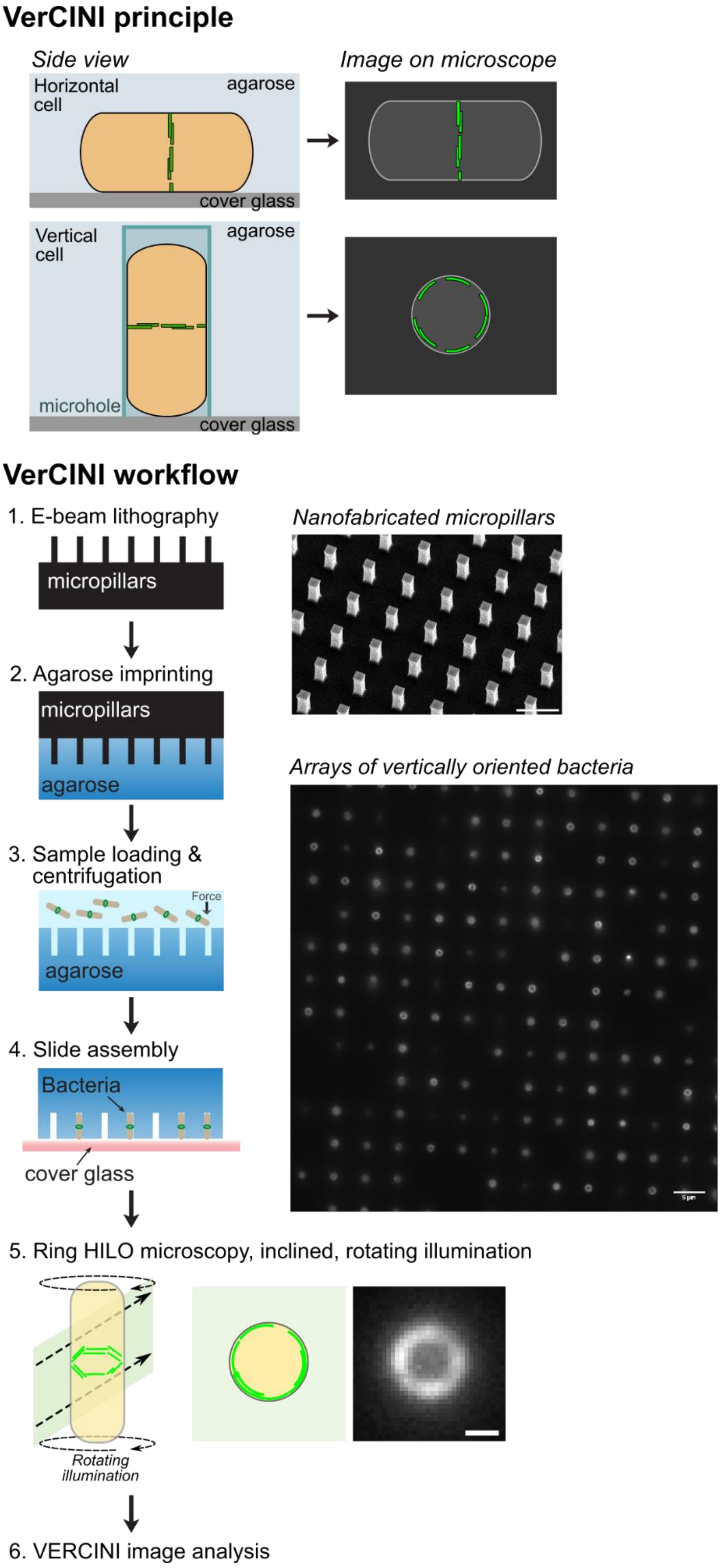


**Supplementary Figure 5: Principle and workflow for Vertical Cell Imaging by Nanostructured Immobilisation (VerCINI).** Nanofabricated cell traps orient bacterial cells vertically on the microscope slide such that the division septum is oriented parallel to the microscope image plane. Silicon micropillars are fabricated by electron-beam lithography (1). Micropillars are used as a mould for molten agarose (2). Cells are loaded into agarose microholes by centrifugation (3) and assembled on a microscope slide (4). Cells are imaged by ring HiLO microscopy for high signal-to-noise ratio and even illumination (5). VerCINI data is then processed using a custom image analysis pipeline (6, Supplementary Figure 4). Micropillar array imaged by Scanning Electron Microscopy. Vertically trapped bacteria expressing FtsZ-GFP (SH130) imaged ring HiLO fluorescence microscopy. Scale bars: 5 µm, top, middle images, 0.5 µm, bottom image.

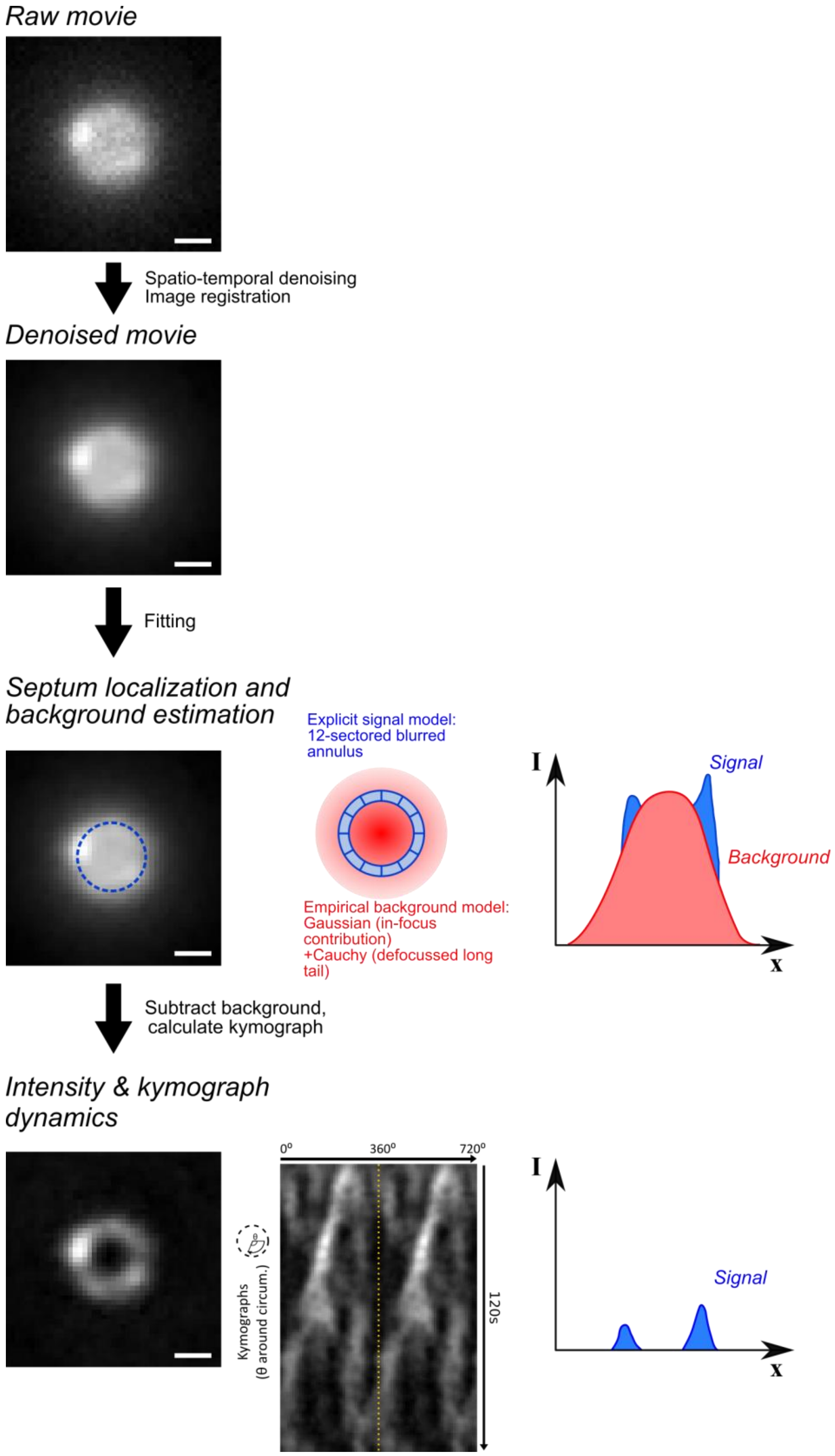


**Supplementary Figure 6: VerCINI image analysis workflow.** Image processing of exemplar VerCINI data for FtsZ-GFP (SH130) strain. VerCINI movie is denoised and registered. Septal localisation is performed by fitting an explicit joint model of a multi-sector blurred annulus for localised signal, plus a mixed Gaussian Cauchy distribution for cytoplasmic background. Fitted background is then subtracted. Intensity around the fitted septal line profile is calculated to sub-pixel accuracy for each movie frame and plotted as a kymograph for further analysis. Scale bars 0.5 µm.

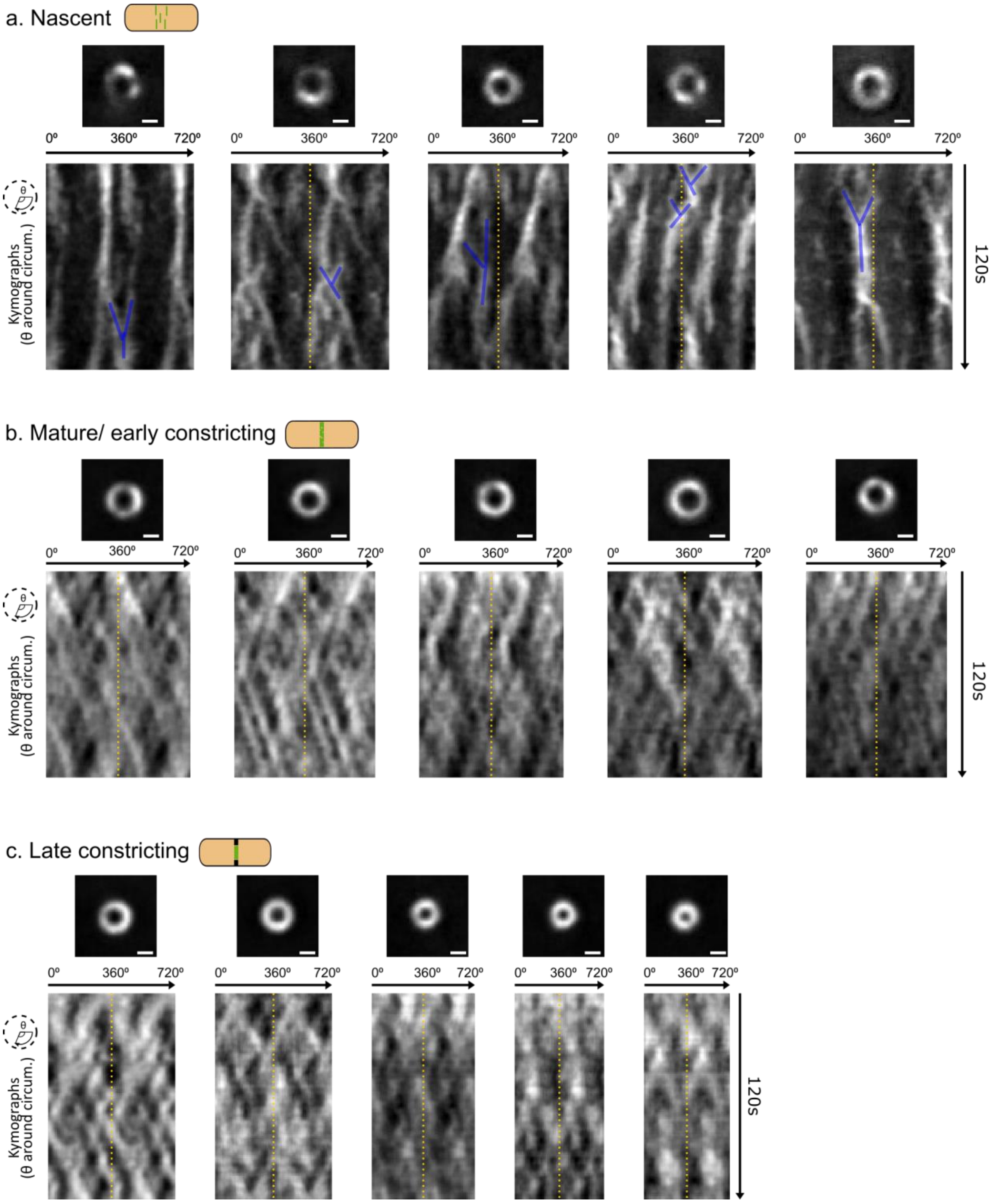


**Supplementary Figure 7: Z-ring dynamics in each cell division phase, nascent (a), mature/ early constricting (b) and late constricting (c) imaged by VerCINI.** Images are snapshots of Z-ring organisation at beginning of time lapse. Kymographs are plotted around Z-ring circumference, with a complete line profile around the circumference repeated twice side-by-side to allow visualisation of filament trajectories which cross the 0º-360º boundary; yellow dashed line indicates beginning of repeated kymograph. Blue lines indicate examples of FtsZ filament interaction. Scale bar 0.5 µm

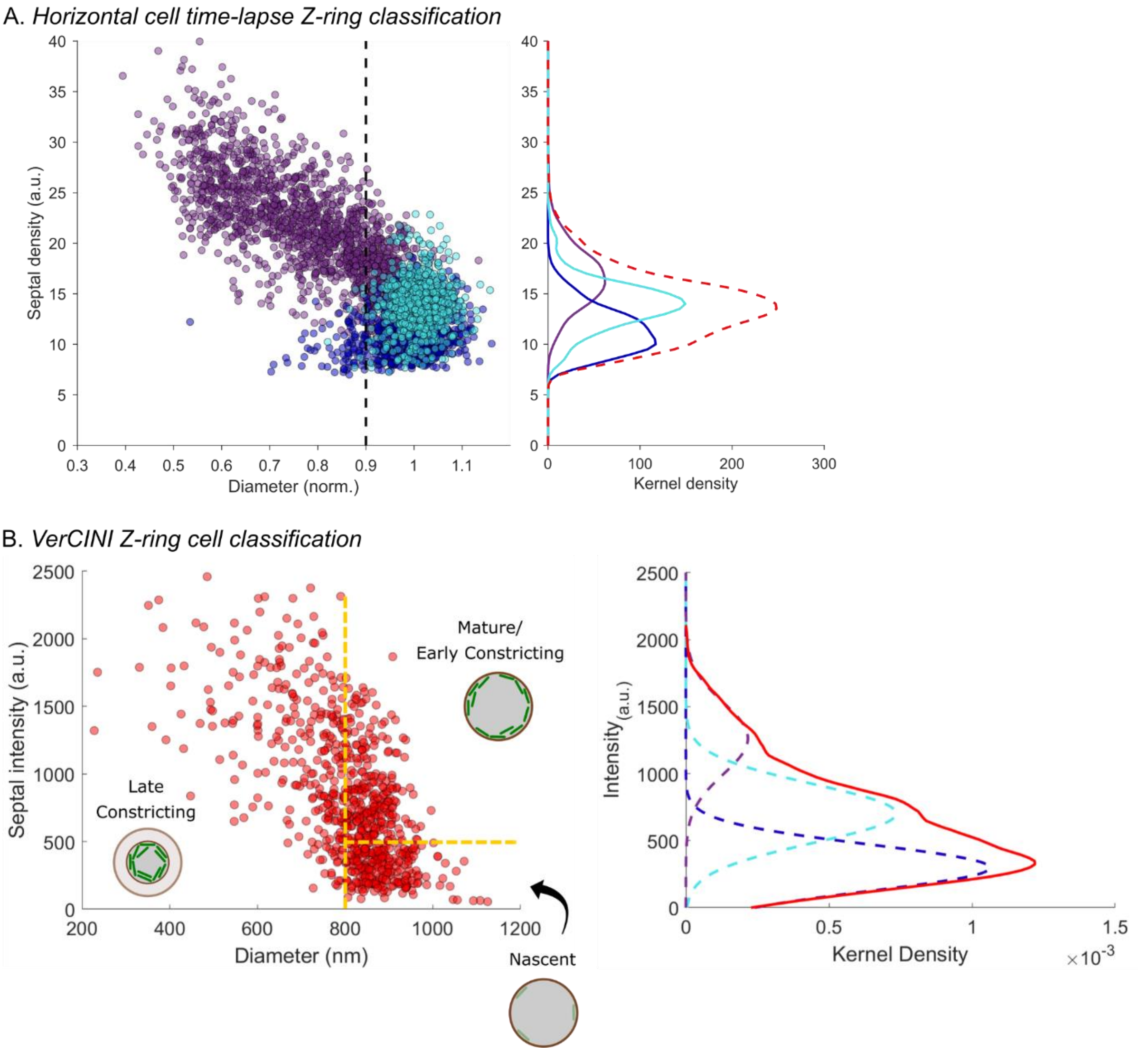


**Supplementary Figure 8: Classification of Z-ring state for horizontal and vertical cells.** (a) Septal density and diameter of Z-rings for time lapse microscopy of horizontal cells. Z-ring phase classified as constricting (purple), mature (cyan) or nascent (magenta) based on time lapse analysis of Z-ring diameter and axial thickness (Figure 1b, Supplementary Note 3). Kernel density plot (right) of septal density for normalized cell diameters > 0.9 (black line, septal density-diameter plot). Continuous lines show kernel density estimates of septal density for each separate Z-ring phase. Red dashed line indicates sum of all 3 distributions. (b) Septal density and diameter of Z-rings for vertical cells imaged by VerCINI. Cells in each identified quadrant were classified as *nascent, mature/ early constricting or late constricting* Z-rings based on septal intensity and diameter, calibrated against horizontal cell time lapse analysis (Supplementary Note 3). Kernel density plot (right, red dashed line) of VerCINI septal intensity for highlighted grey region of septal density-diameter plot. Dashed lines show contribution of nascent (blue), mature (cyan) and constricting (purple) to observed septal intensity distribution, as estimated by fitting a 3 Gaussian model to the data.

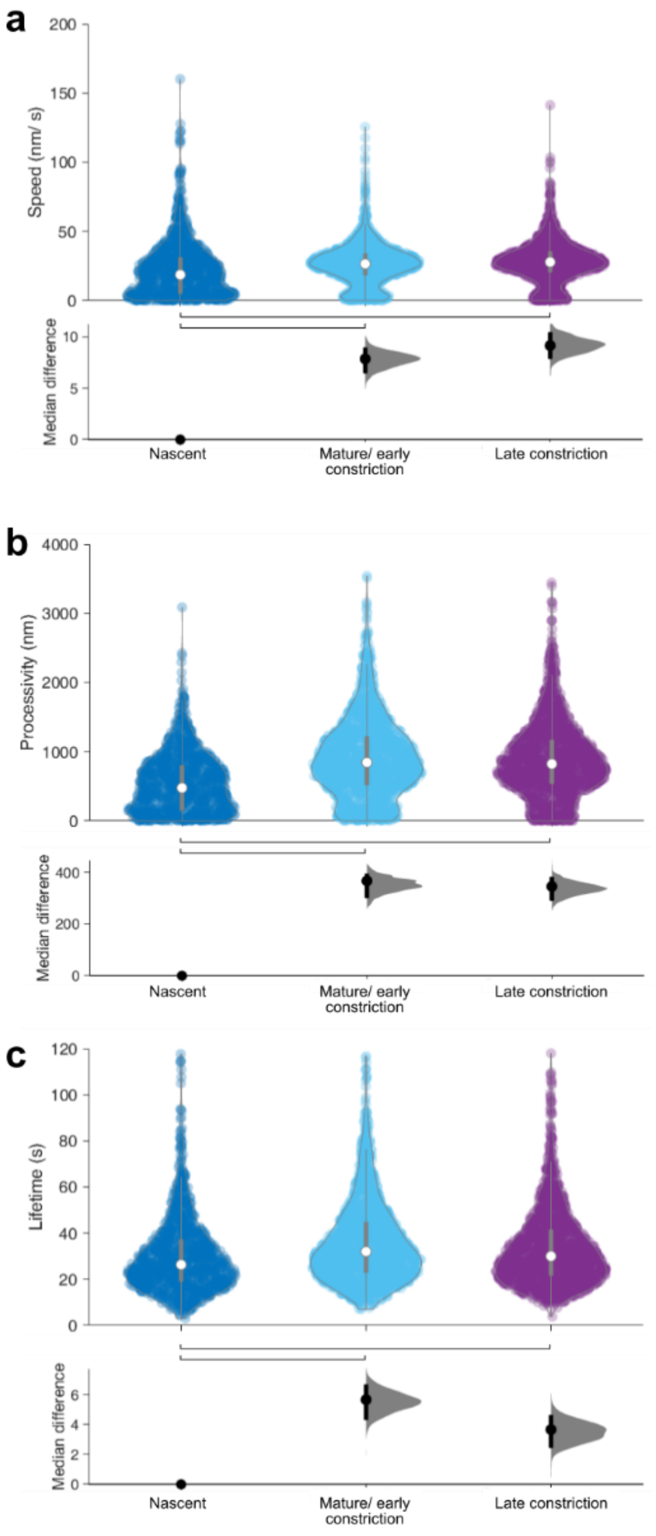


**Supplementary Figure 9: Quantification of FtsZ-GFP (SH130) filament dynamics for cells imaged by VerCINI, with DABEST analysis of median difference for each cell cycle stage**. (a) Speed distribution of filaments (as presented Figure 1h, with additional DABEST analysis). (b) Filament processivity (distance filament treadmills around septal circumference). (c) Filament lifetime (time bound to septum). Violin plot: white circles, median; thick grey lines, interquartile range; thin grey lines, 1.5x interquartile range. DABEST plot: black circle, median difference between indicated conditions; black lines, 95% confidence interval of median difference.

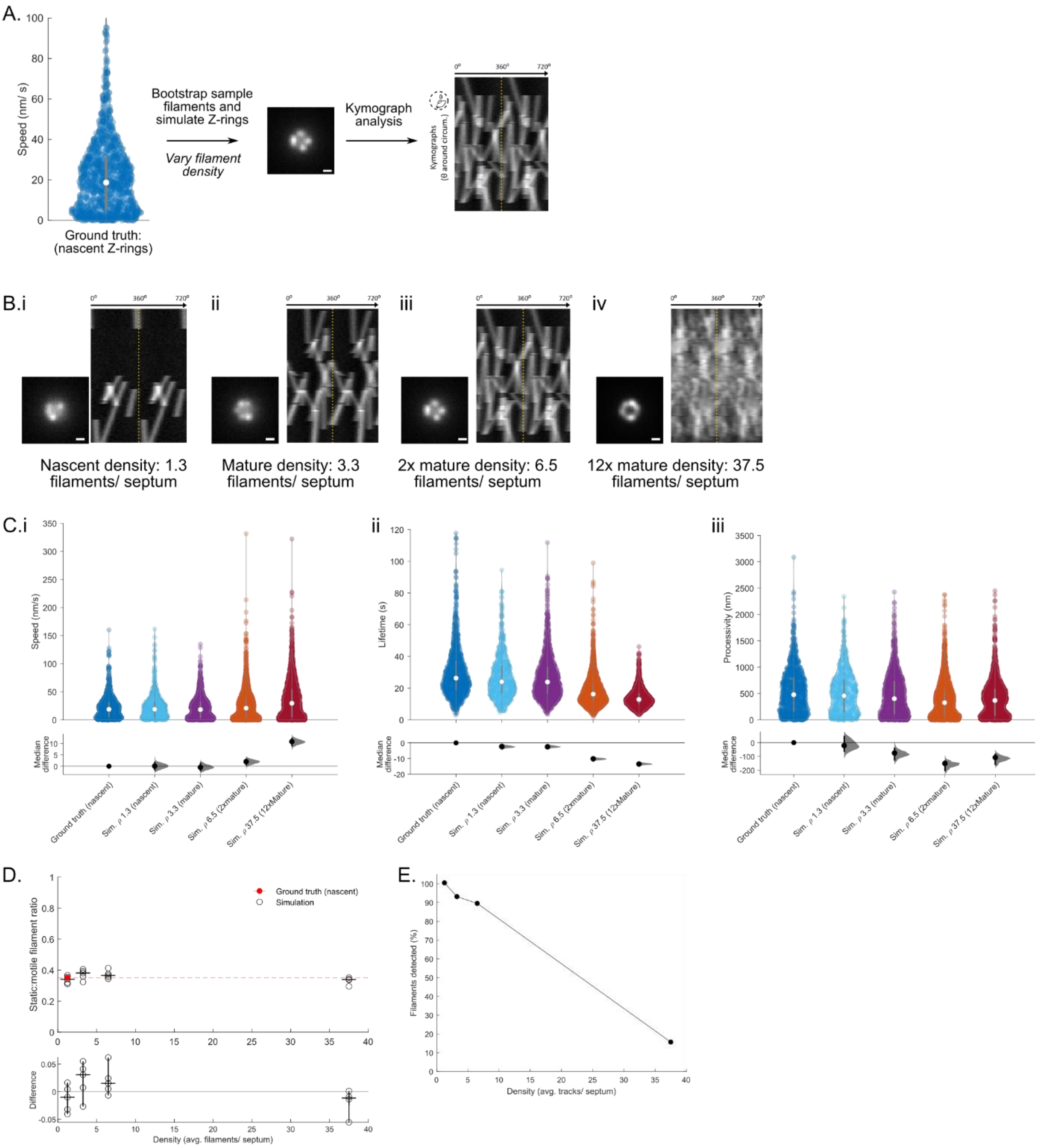


**Supplementary Figure 10: Validation of kymograph filament speed analysis workflow on simulated data.** (a) Simulation workflow: Simulated filament speeds are bootstrap sampled from observed experimental FtsZ filament speed distribution in nascent Z-rings (also presented in Figure 1h). FtsZ rings are simulated per Methods. Kymograph analysis is performed on the simulated data. (b) Exemplar Z-ring images and kymographs of simulated data. (c) Observed filament speed, lifetime and processivity distribution for ground truth and simulated at indicated filament density, with DABEST analysis. Violin plot: white circles, median; thick grey lines, interquartile range; thin grey lines, 1.5x interquartile range. DABEST plot: black circle, median difference between indicated conditions; black lines, 95% confidence interval of median difference. (d) Estimated fraction of static filaments at each simulated density (hollow black circles) compared to ground truth static filament fraction (filled red circles). Each hollow circle, result for 20 simulated Z-rings. Bottom panel, difference between observed static filament fraction and ground truth. Horizontal line, median. Error bars, 95% CI. (e) Percentage of simulated filaments detected at each simulated density. Scale bars, 0.5 µm.

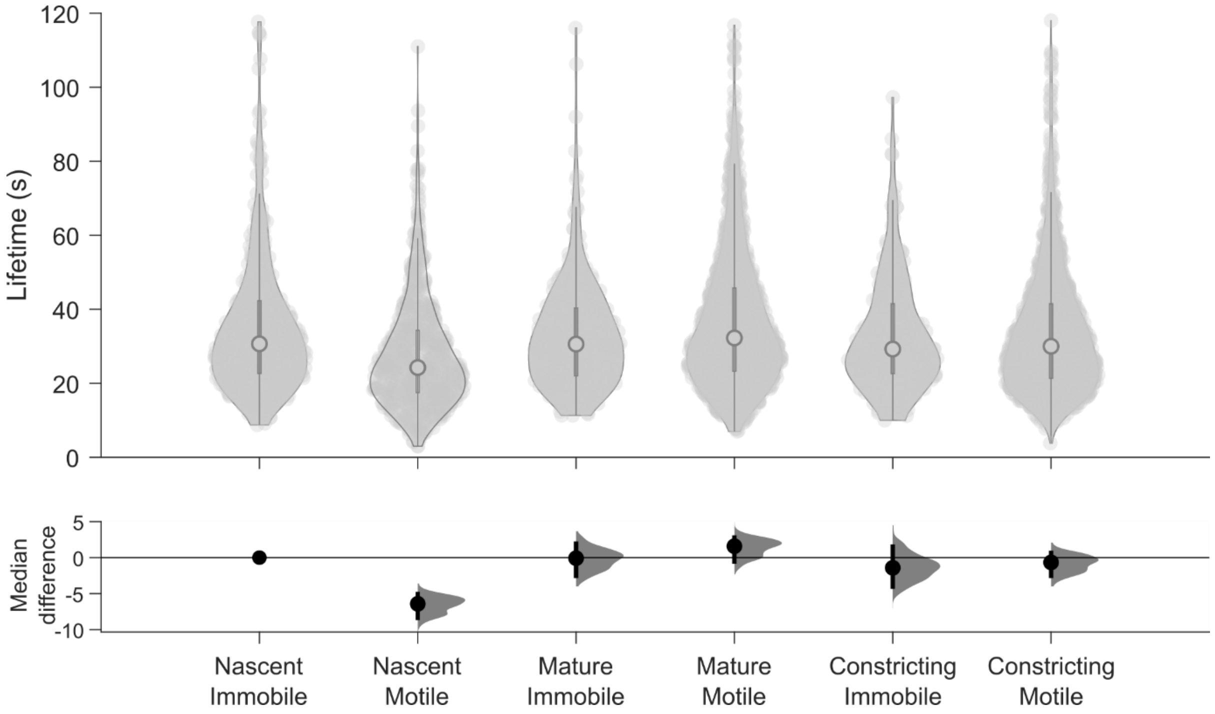


**Supplementary Figure 11: Quantification of FtsZ-GFP (SH130) filament binding lifetime for cells imaged by VerCINI, with DABEST analysis of median difference for each cell cycle stage, classified by filament motility (motile/ immobile, filament speed threshold 10 nm/ s)**. Violin plot: white circles, median; thick grey lines, interquartile range; thin grey lines, 1.5x interquartile range. DABEST plot: black circle, median difference between indicated conditions; black lines, 95% confidence interval of median difference.

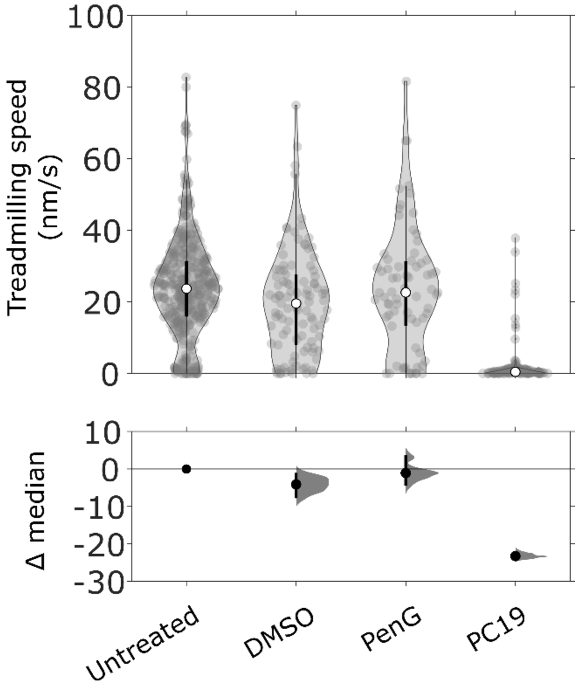


**Supplementary Figure 12: Quantification of FtsZ-GFP (SH130) filament dynamics during chemical perturbations with DABEST analysis.** Violin plot: white circles, median; thick black lines, interquartile range; thin black lines, 1.5x interquartile range. DABEST plot: black circle, median difference between indicated conditions; black lines, 95% confidence interval of median difference.

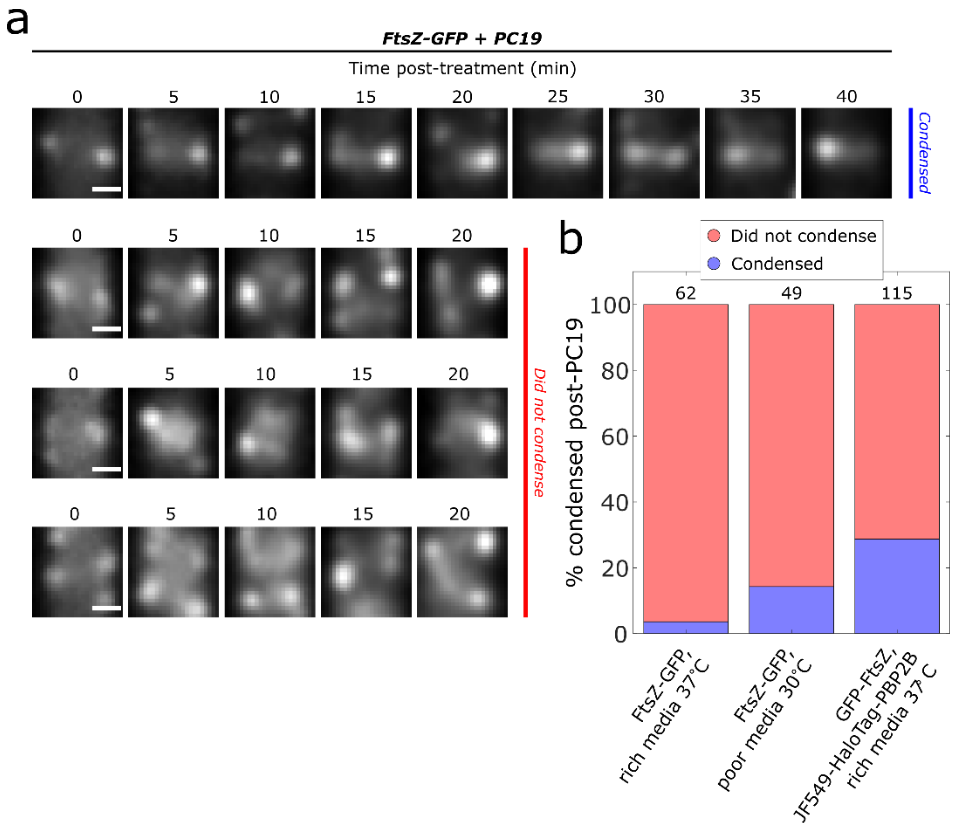


**Supplementary Figure 13: Condensation of Z-rings post-PC19 across strains and conditions.** (a) Representative time-lapses of nascent Z-rings for FtsZ-GFP cells (SH130) after arrival of 10 µM PC19-laced media. Nascent Z-rings typically fail to condense into mature Z-rings post-treatment. Scale bars: 500 nm. (b) Stacked bar plot showing the percentage of cells in the nascent Z-ring stage at t=0 min that condensed post-PC19 treatment under three conditions (FtsZ-GFP strain SH130; GFP-FtsZ, HaloTag-PBP2B strain SH212). Z-rings classified as nascent based on thresholds: thickness>400 nm (SH130) or thickness>450 nm (SH212). Larger threshold for SH212 strain is due to poor SNR (see Supplementary Figure 11). Numbers above stacked bars indicate number of cells. The higher percentage of SH212 cells condensed likely results from lower SNR in these experiments leading to misclassification of some already-condensed, mature Z-rings as 'nascent'.

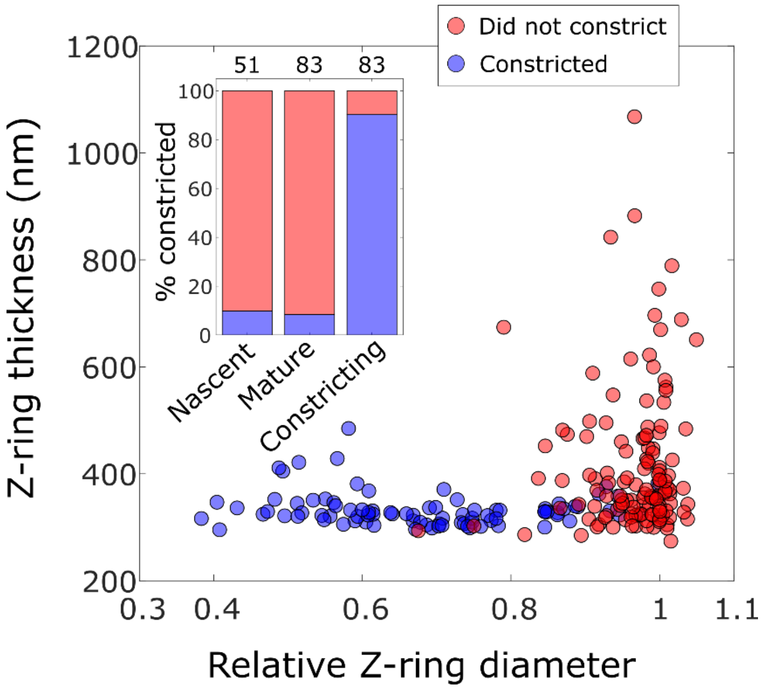


**Supplementary Figure 14: Septal completion assay post-PC19 treatment, slow growth conditions (minimal media, 30°C).** FtsZ-GFP strain SH130. Scatter plot of Z-ring diameters and thicknesses for all FtsZ-GFP cells at t=0 min showing whether they constricted (blue) or not (red). *Inset*: Percentage of cells that constricted classified by division stage. Nascent: thickness>400 nm, mature: thickness<400 nm, relative diameter>0.9, constricting: thickness<400 nm, relative diameter<0.9. Numbers above stacked bars indicate number of cells.

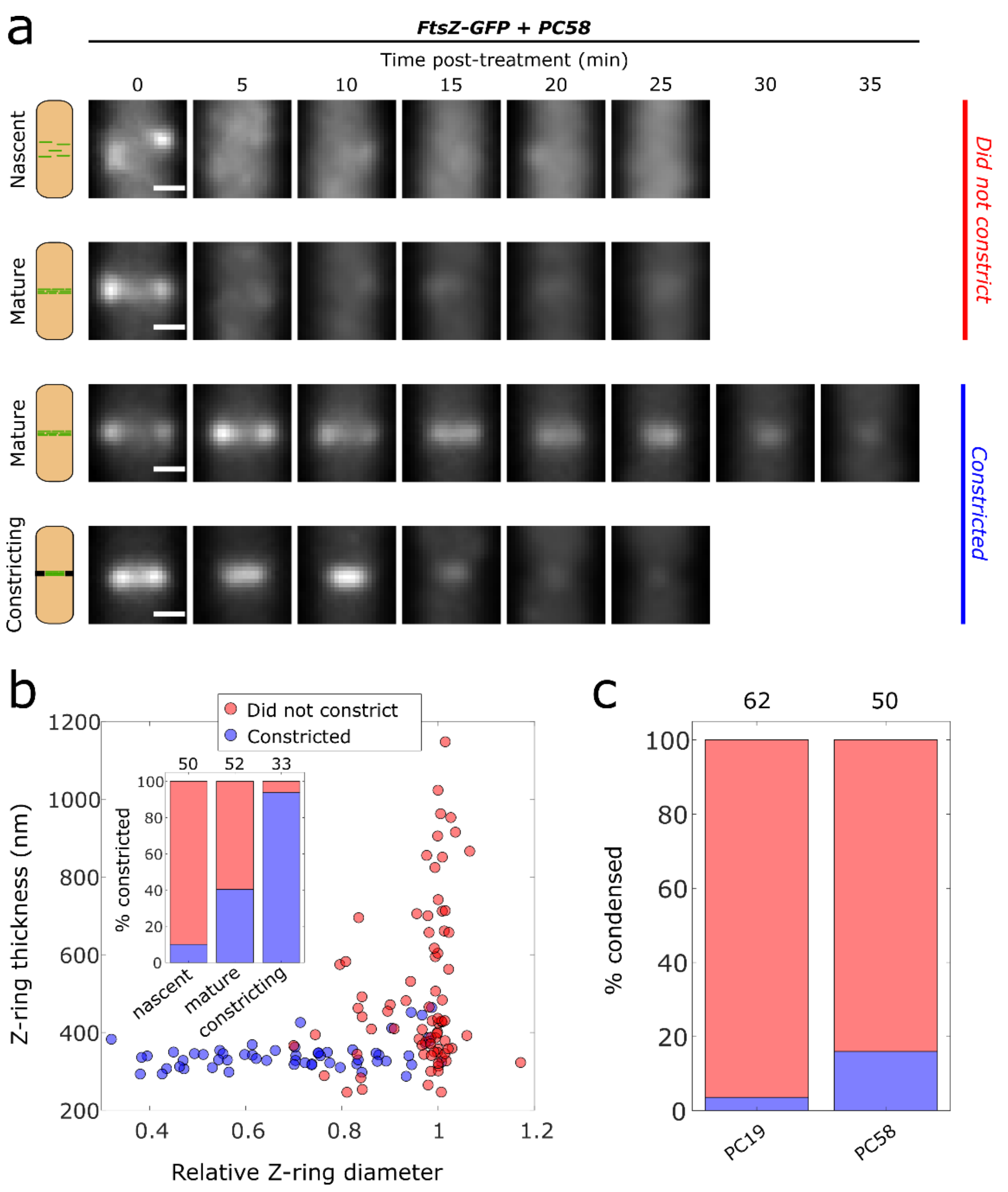


**Supplementary Figure 15: Septal completion assay post-PC58 treatment.** (a) Representative time-lapses of Z-rings for FtsZ-GFP cells (SH130) after arrival of 1 mM PC58-laced media. Most nascent rings and many mature rings do not constrict for tens of minutes after PC58 treatment, whereas constricting rings and many mature rings continue constricting. Scale bars: 500 nm. (b) Scatter plot of Z-ring diameters (normalized by maximal diameter for each Z-ring) and thicknesses for all FtsZ-GFP cells at t=0 min showing whether they continued constricting (blue) or not (red). *Inset*: Percentage of cells that continued constricting classified by division stage. Nascent: thickness>400 nm, mature: thickness<400 nm, relative diameter>0.9, constricting: thickness<400 nm, relative diameter<0.9. (c) Percentage of cells in the nascent Z-ring stage that condensed post-PC58 compared to the percentage that condensed post-PC19. Numbers above stacked bars indicate number of cells.

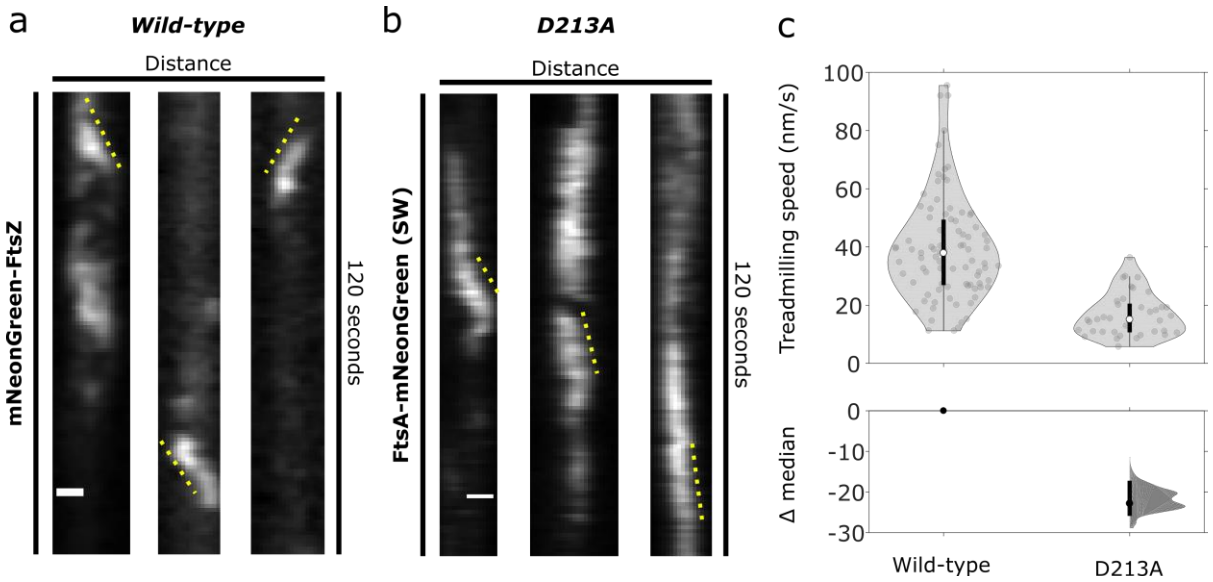


**Supplementary Figure 16: Effect of FtsZ(D213A) expression on treadmilling speed.** Representative kymographs of filament treadmilling dynamics measured by TIRF microscopy for mNeonGreen-FtsZ (strain bWM4) (a) and FtsA-mNeonGreen under dominant negative expression of FtsZ(D213A) (strain bAB217) (b) in rich media 37°C. bAB217 cells were induced with 10 µM IPTG induction for 1 hour prior to imaging. Filament speed is determined by manually tracing filament trajectories on the kymograph (indicated by yellow dashed line) and calculating the angle of the traced line. Scale bar: 500 nm. (c) Distribution of filament treadmilling speeds for both conditions. Violin plot: white circles, median; thick black lines, interquartile range; thin grey lines, 1.5x interquartile range. DABEST plot: black circle, median difference between indicated conditions; black lines, 95% confidence interval of median difference.

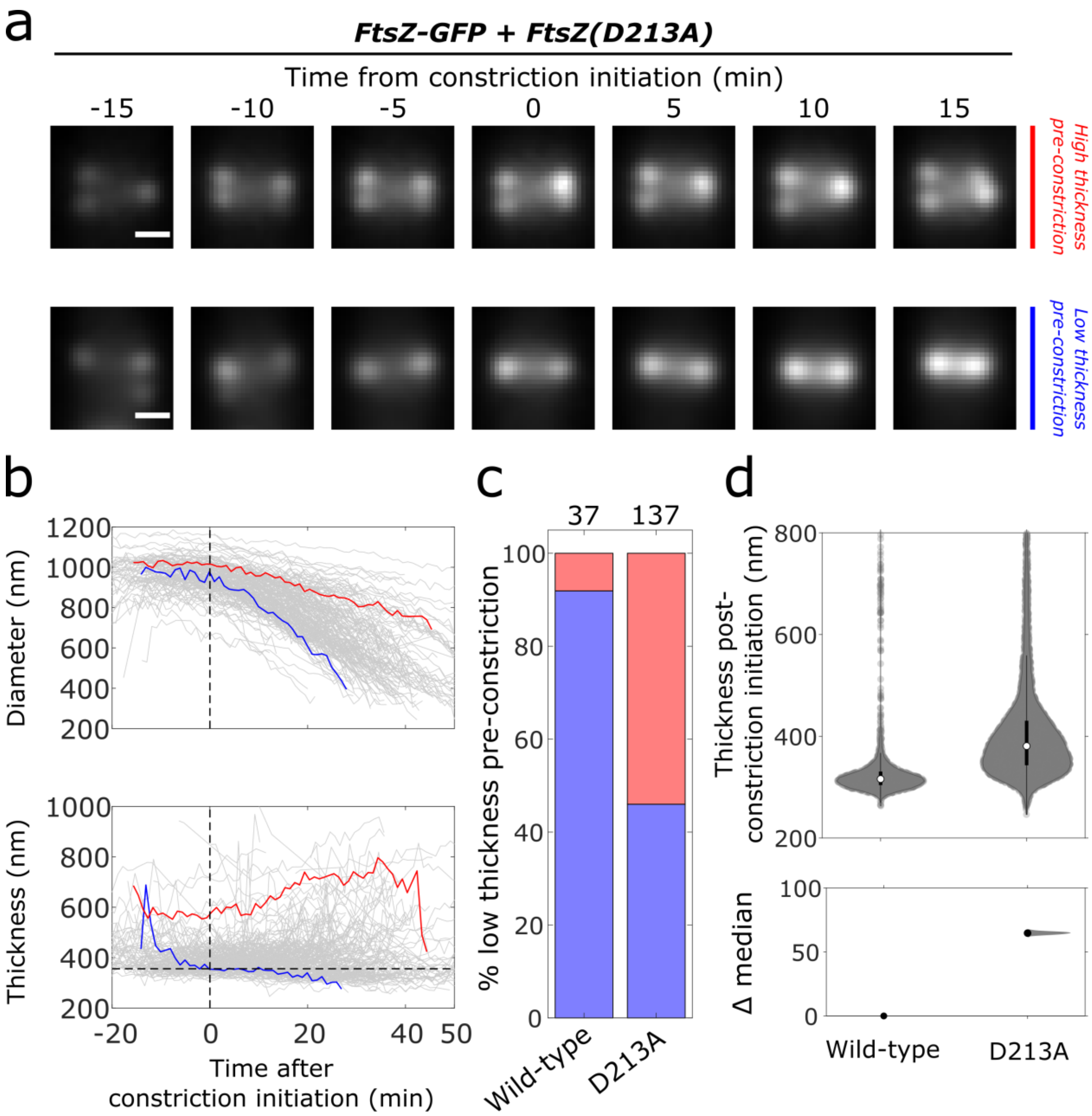


**Supplementary Figure 17: Effect of FtsZ(D213A) expression on Z-ring condensation.** (a) Representative time-lapses of Z-rings in a strain expressing FtsZ-GFP from the native locus and the GTPase-deficient mutant FtsZ(D213A) from an inducible promoter (strain SH131 induced with 10 µM IPTG for 1 hour prior to imaging). t=0 represents the start time of constriction as determined from fits to constriction trajectories. Scale bars: 500 nm. (b) Trajectories of Z-ring diameter and thickness over time aligned to constriction start time at t=0. Highlighted trajectories are those of the representative traces in (a) (Red: high thickness pre-constriction, Blue: low thickness pre-constriction). Vertical dotted lines: start time of constriction (t=0). Horizontal dotted line: cutoff for determining ‘high thickness’ versus ‘low thickness’ (mean + 1σ from distribution of condensed wild-type cells; 356 nm). (c) Percentage of Z-rings that had low thickness prior to the start of constriction for wild-type (strain SH130) and 10 µM IPTG induction of D213A (SH131 strain). Numbers above stacked bars indicate number of cells. (d) Thickness of Z-rings after start of constriction for wild-type and 10 µM IPTG induction of D213A. Violin plot: white circles, median; thick black lines, interquartile range; thin black lines, 1.5x interquartile range. DABEST plot: black circle, median difference between indicated conditions; black lines, 95% confidence interval of median difference.

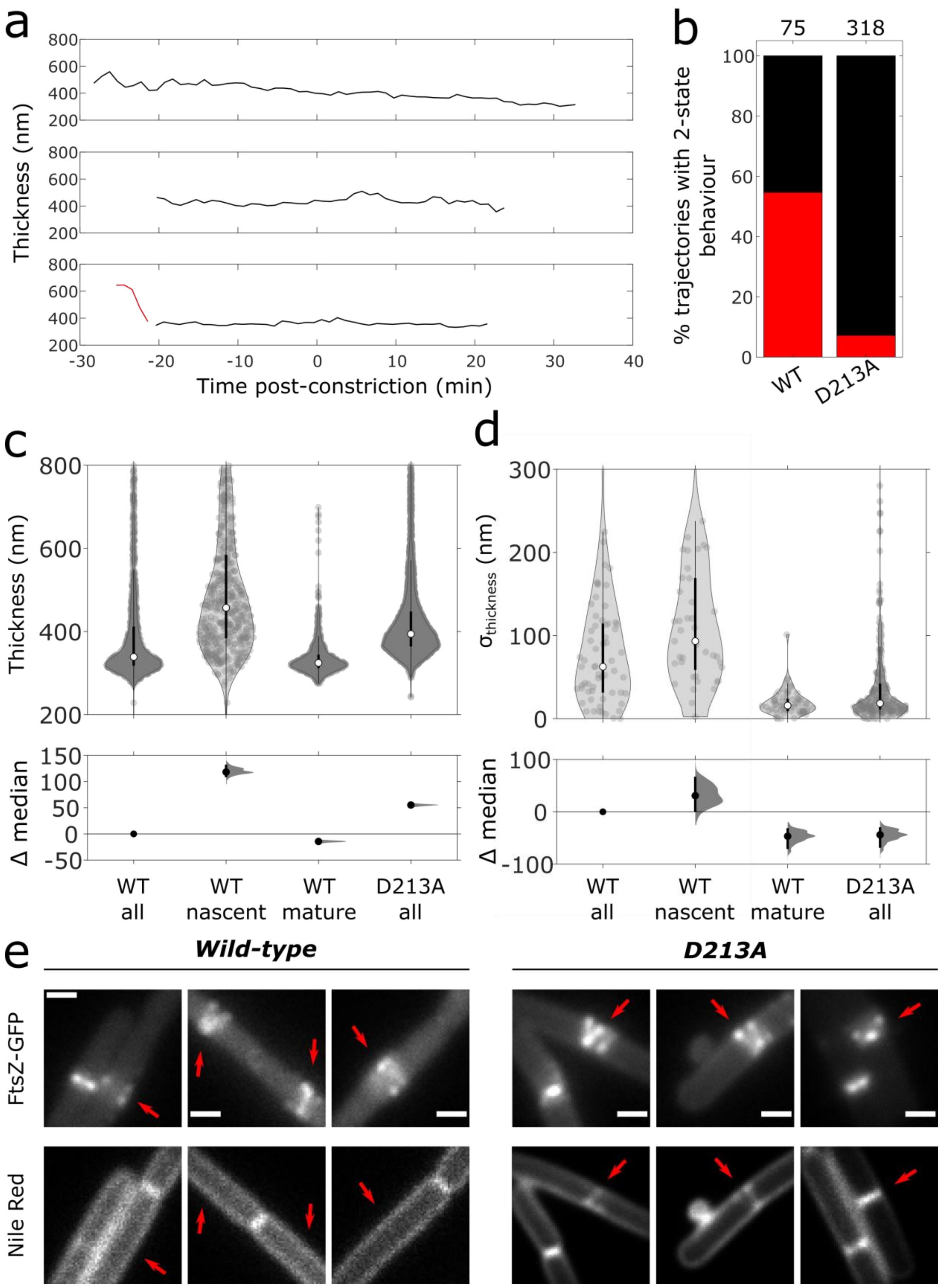


**Supplementary Figure 18: Comparison of pre-constriction Z-rings between wild-type (SH130) and FtsZ(D213A) mutant (SH131).** (a) Exemplar traces of classified time series of Z-ring axial thickness for SH131 cells. Red line indicates decondensed state assigned by changepoint detection, black line indicates condensed state. (b) Percentage of trajectories showing two states prior to the start of constriction for wild-type (strain SH130; 55%) and 10 µM IPTG induction of D213A (SH131 strain; 7%). Numbers above stacked bars indicate number of trajectories. (c and d) Z-ring thickness (c) and standard deviations of Z-ring thickness (d) pre-constriction for wild-type and 10 µM IPTG induction of D213A. Distributions for wild-type are separated into nascent and mature Z-ring states by state change identification (Methods). Violin plot: white circles, median; thick black lines, interquartile range; thin black lines, 1.5x interquartile range. DABEST plot: black circle, median difference between indicated conditions; black lines, 95% confidence interval of median difference. (e) Representative images of Z-rings with large thicknesses in a strain expressing FtsZ-GFP from the native locus (SH130 strain; "wild-type" in figure) and a strain expressing both FtsZ-GFP from the native locus and the GTPase-deficient mutant FtsZ(D213A) from an inducible promoter (SH131 strain induced with 10 µM IPTG for 1 hour prior to imaging; "D213A" in figure) with corresponding membrane stain (Nile Red). Top row: FtsZ-GFP signal. Bottom row: Nile Red signal. Red arrows point to Z-rings and corresponding membrane invaginations where present. For visibility, Nile Red images use gamma correction: wild-type images use gamma=2, D213A images use gamma=4. Scale bars: 1 µm.

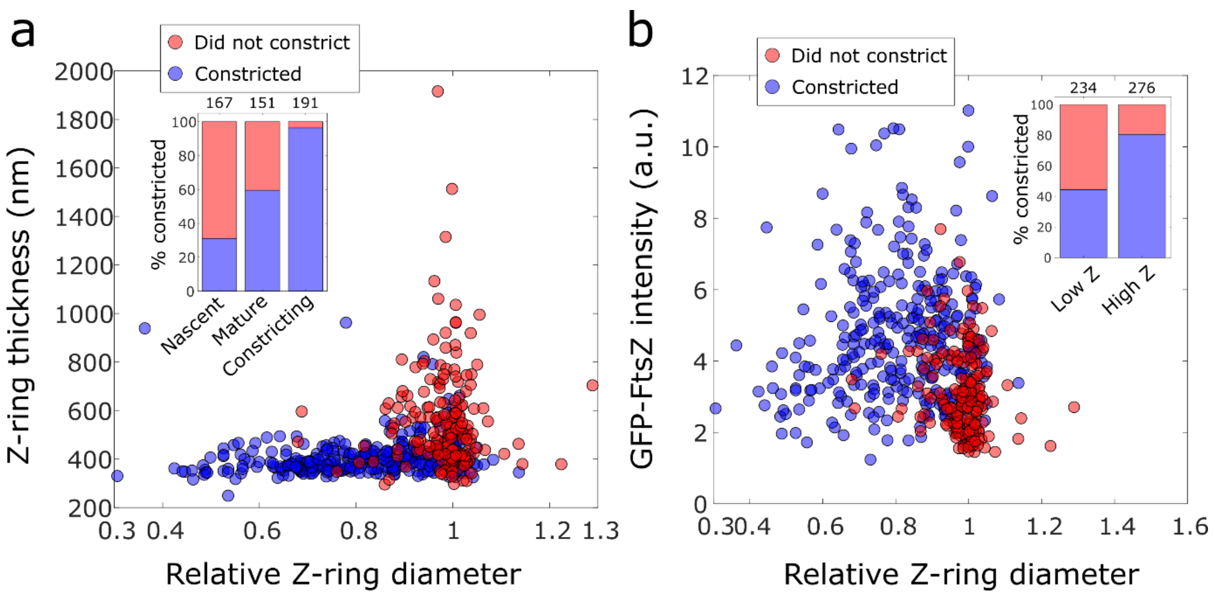


**Supplementary Figure 19: Septal constriction assay post-PC19 treatment, GFP-FtsZ HaloTag-PBP2B strain (SH212).** (a) Scatter plot of Z-ring diameters and thicknesses for all GFP-FtsZ HaloTag-PBP2B cells labelled with JF549 HaloTag ligand at t=0 min showing whether they constricted (blue) or not (red). *Inset*: Percentage of cells that constricted classified by division stage. Nascent: thickness>450 nm, mature: thickness<450 nm, relative diameter>0.9, constricting: thickness<450 nm, relative diameter<0.9. (b) Scatter plot of Z-ring diameters and intensities of GFP-FtsZ signal at t=0 min showing whether they constricted (blue) or not (red) for mature rings only. *Inset*: Percentage of cells that constricted classified by intensity of GFP-FtsZ. Low Z: intensity<1.2, high Z: intensity>1.2. Numbers above stacked bars indicate number of cells.

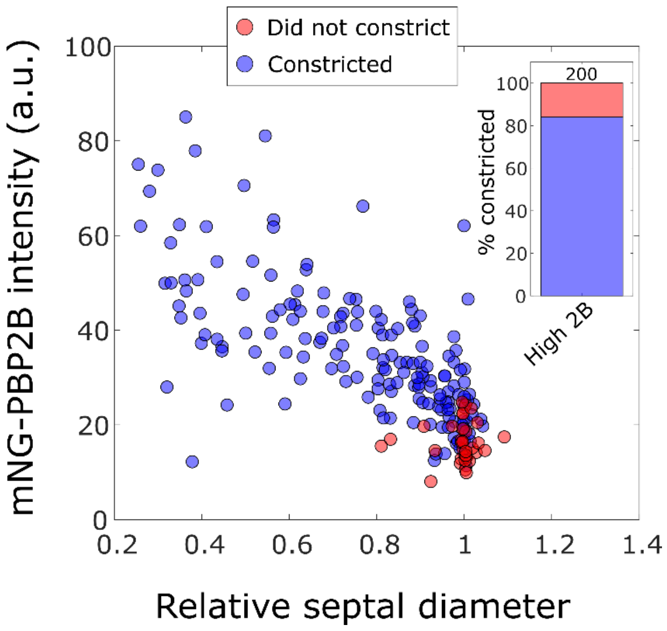


**Supplementary Figure 20: Septal constriction assay post-PC19 treatment, mNeonGreen-PBP2B strain (ME7).** Scatter plot of septal diameters and intensities of mNeonGreen-PBP2B signal at t=0 min showing whether they constricted (blue) or not (red) for mature rings only. *Inset*: Percentage of cells that constricted. Number above stacked bar indicates number of cells.

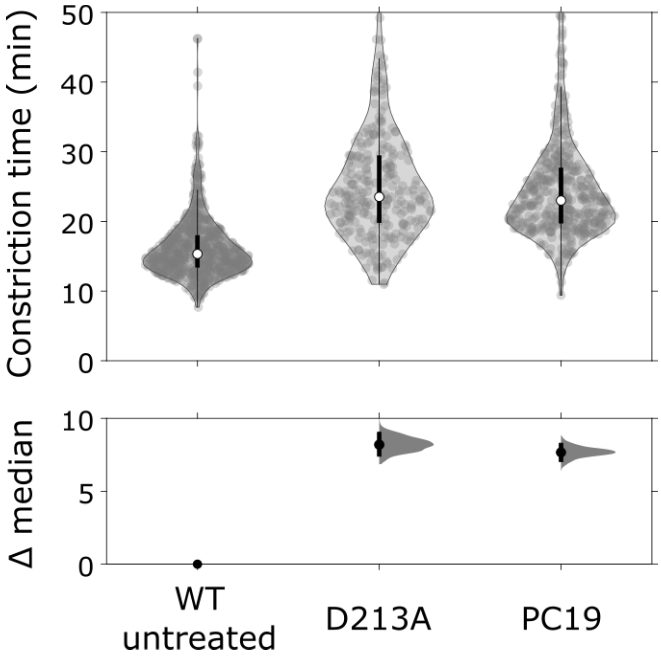


**Supplementary Figure 21: Effect of high D213A expression on constriction time.** *Top panels:* Violin plots of constriction time comparing wild-type untreated cells (ME7 strain) to cells expressing high levels of the GTPase-deficient mutant FtsZ(D213A) (strain SH132 induced with 10 µM IPTG for 1 hour prior to imaging) and wild-type cells treated with PC19 (ME7 strain). White circles, median; thick black lines, interquartile range; thin black lines, 1.5x interquartile range. *Bottom panels:* DABEST plots showing the effect sizes on constriction time. Black circle, median difference between indicated conditions; black lines, 95% confidence interval of median difference.

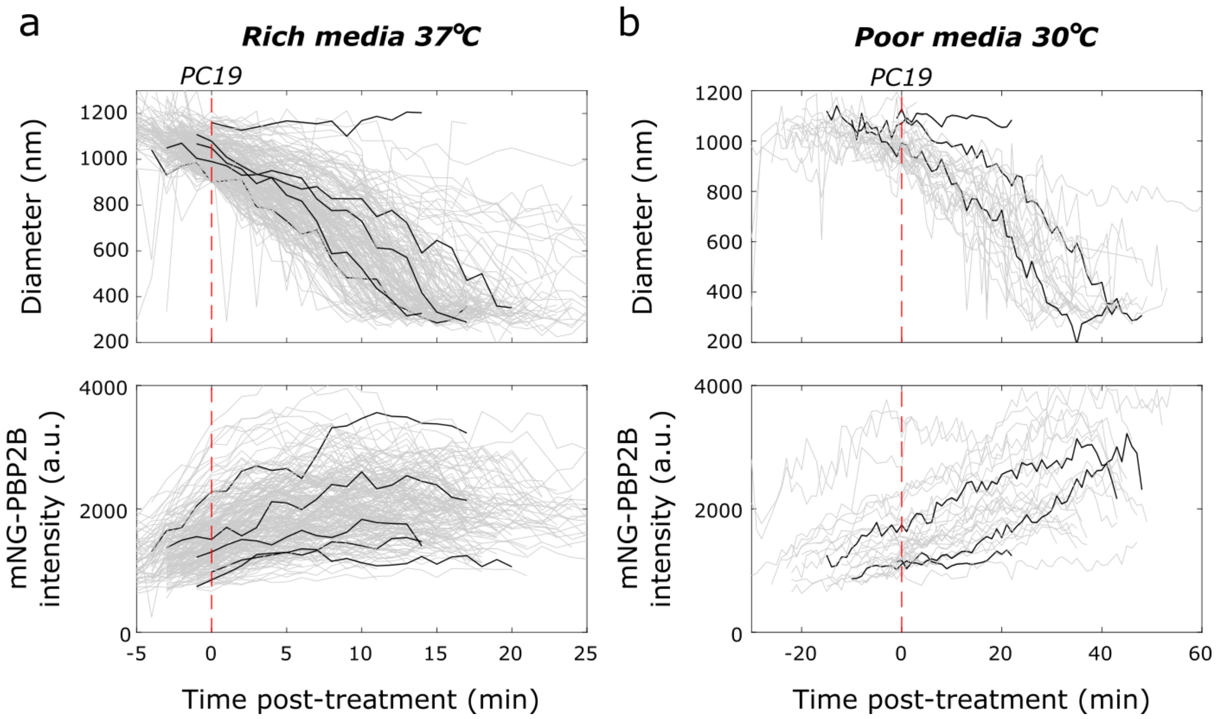


**Supplementary Figure 22: Intensity of mNeonGreen-PBP2B (ME7) post-PC19 treatment.** *Grey:* All traces for this condition. *Black:* Representative traces. *Red dotted lines:* Time of PC19 treatment. (a) Diameters and intensities of mNeonGreen-PBP2B cells in fast growth conditions (rich media 37°C) (b) Diameters and intensities in slow growth conditions (poor media 30°C).

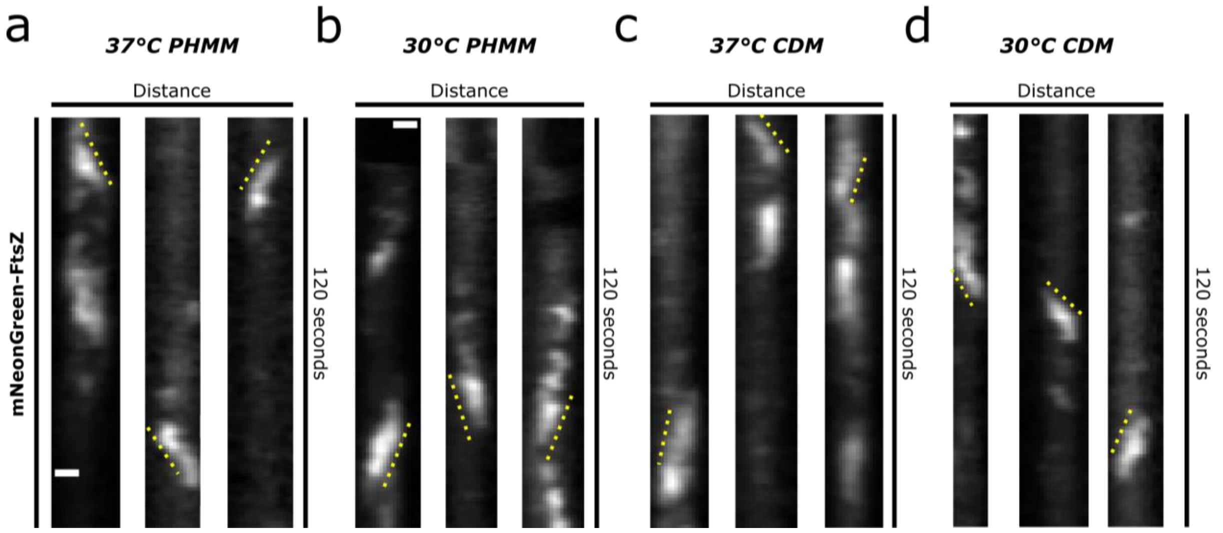


**Supplementary Figure 23: Representative kymographs of filament treadmilling dynamics measured by TIRF microscopy for mNeonGreen-FtsZ (strain bWM4) under different growth conditions.** Filament speed is determined by manually tracing filament trajectories on the kymograph (indicated by yellow dashed line) and calculating the angle of the traced line. Scale bars: 500 nm.

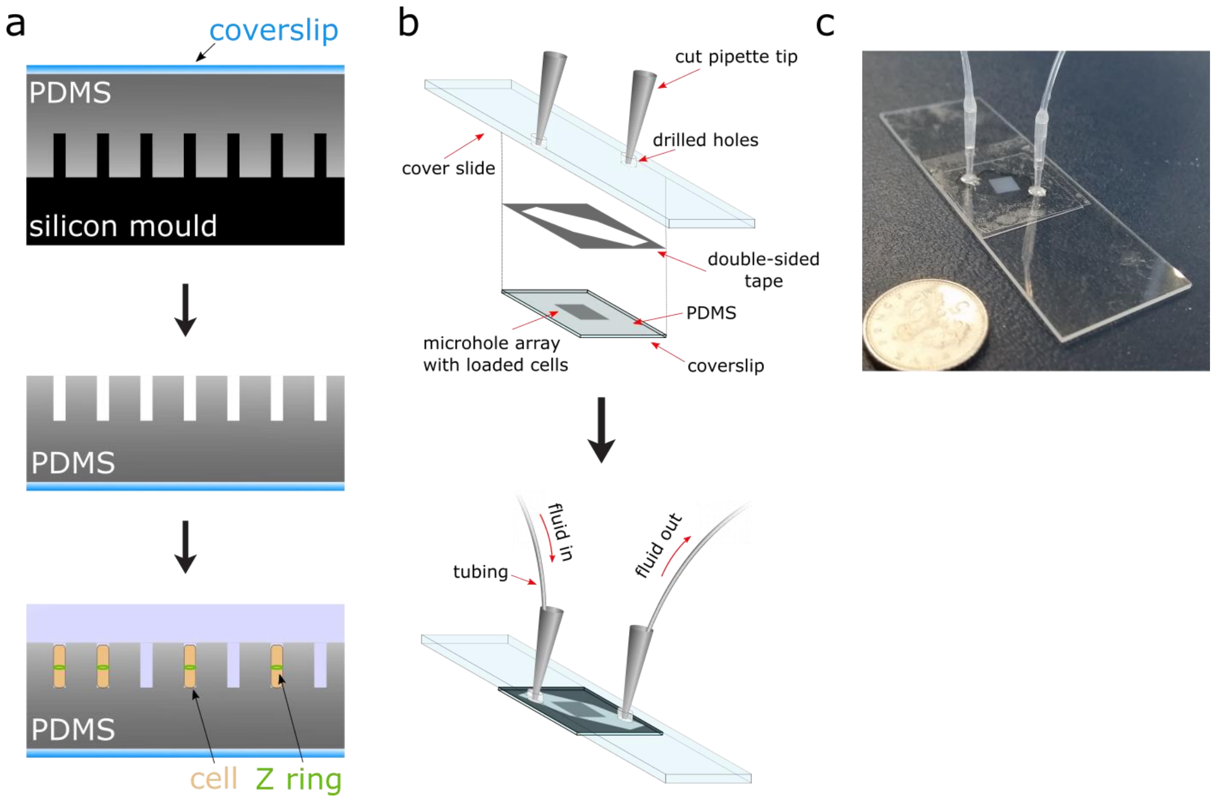


**Supplementary Figure 24: Assembly and preparation of microfluidic VerCINI**. (a) Formation and loading of open-topped microholes. *Top:* PDMS is poured onto a silicon mould containing micropillars and pressed down with a microscope coverslip to form a thin layer. *Middle:* After curing, the PDMS and coverslip are peeled off the silicon mould. *Bottom:* Cells are loaded into the open-topped holes by centrifugation. (b) Assembly of the microfluidic chamber. *Top:* Exploded diagram of fluidic chamber. A cover slide with drilled holes has cut pipette tips inserted and epoxied in place. A piece of double-sided tape has a groove cut into it to serve as a flow channel. This double-sided tape seals the cover slide to the PDMS-coated coverslip containing cells loaded into microholes. *Bottom:* Assembled fluidic chamber. Tubing is inserted into the pipette tips, and fluid flows through the channel formed by the double-sided tape. (c) Photo of assembled chamber with 5p coin for scale.

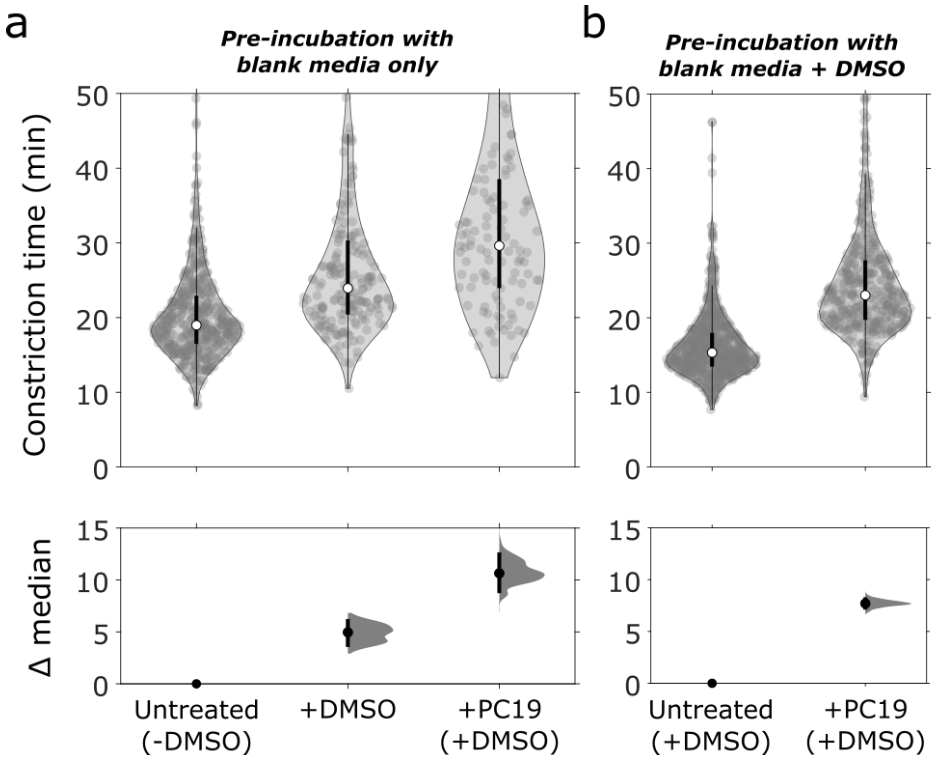


**Supplementary Figure 25: Effect of DMSO on mNeonGreen-PBP2B (ME7) constriction time.** *Top panels:* Violin plots of constriction time. White circles, median; thick black lines, interquartile range; thin black lines, 1.5x interquartile range. *Bottom panels:* DABEST plots showing the effect sizes on constriction time. Black circle, median difference between indicated conditions; black lines, 95% confidence interval of median difference. (a) Constriction times for cells pre-incubated in blank media (with no DMSO) before and after treatment with 1% DMSO or PC19 (which contained 1% DMSO as accompanying solvent). (b) Constriction times for cells pre-incubated in blank media supplemented with 1% DMSO before and after treatment with PC19. Pre-incubation with 1% DMSO ensured that the concentration of DMSO remained constant during PC19 treatment.

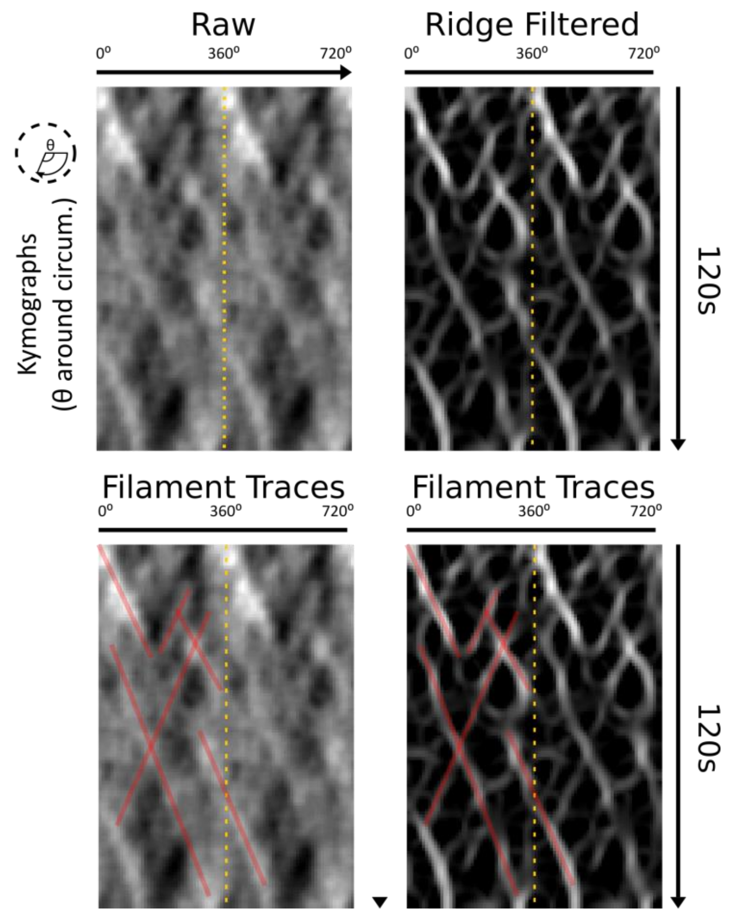


**Supplementary Figure 26: VerCINI FtsZ treadmilling dynamics analysis workflow**. Ridge filter is applied to raw kymographs to detect filament trajectories independent of intensity. Filament trajectories are then manually traced with reference to both the raw and ridge-filtered kymographs; only filament trajectories clearly identifiable in both images were selected. Filament trajectories are exported, and filament speed, processivity and lifetime are calculated. Two full revolutions around the cell (0-720°) are plotted side-by-side in each kymograph to better show continuity of filament tracks as they pass 0°/360°, separated by yellow dotted lines. Red lines indicate all identified trajectories, shown only in the first of the two side-by-side kymographs.

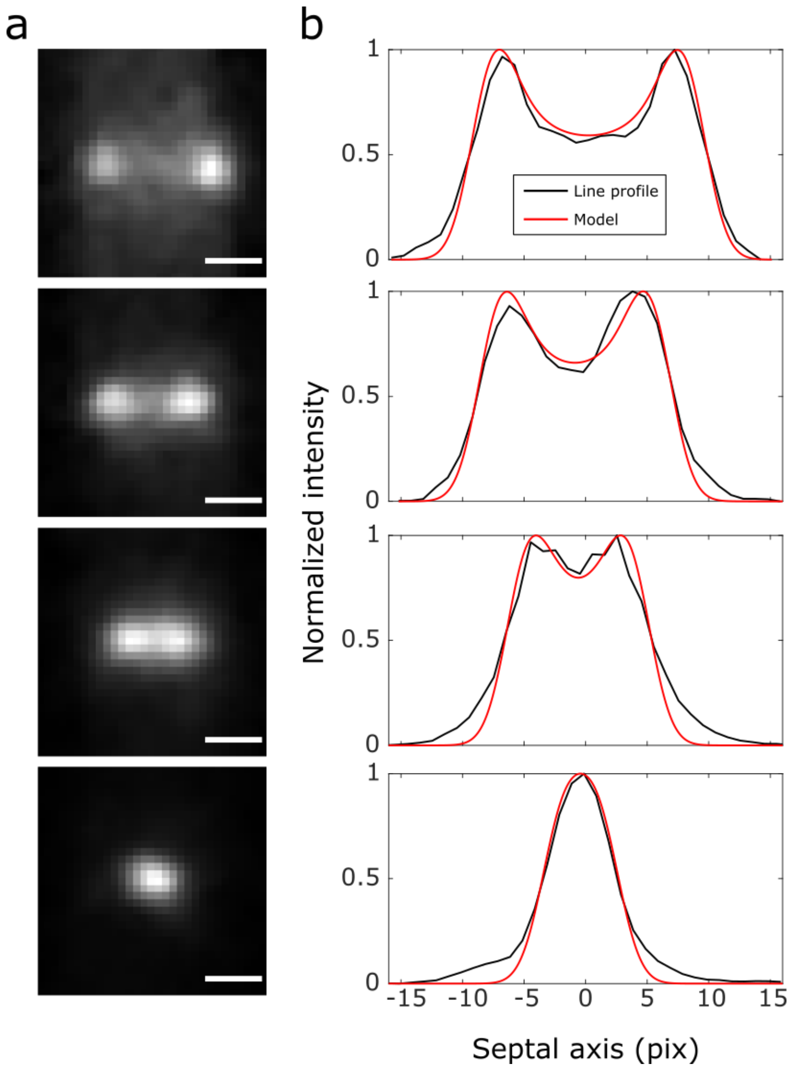


**Supplementary Figure 27: Example of fitting horizontal ring line profile to tilted circle model.** (a) Representative image sequence of a constricting ring of mNeonGreen-PBP2B (ME7). Scale bars: 500 nm. (b) Corresponding line profiles for the images in (a) (black) with fitted tilted circle model (red; Supplementary Note 1).

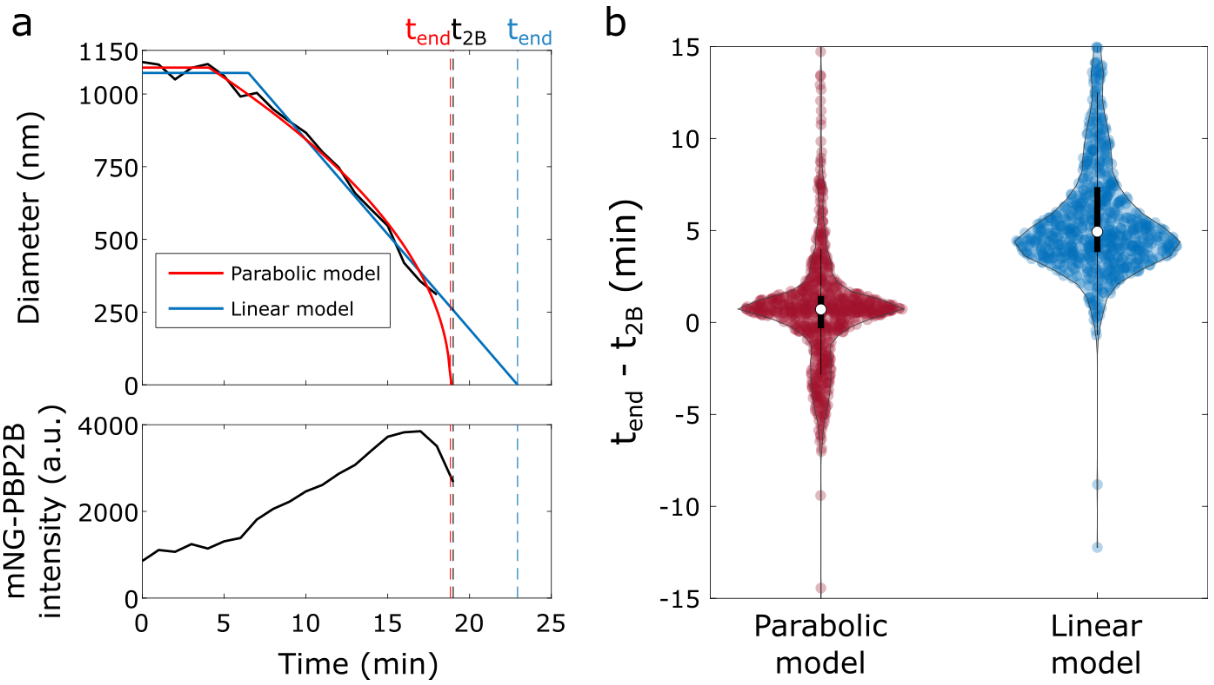


**Supplementary Figure 28: Comparison of linear, parabolic constant synthesis constriction models.** (a) Representative trace of an mNeonGreen-PBP2B (ME7) cell constricting with fitted parabolic (red) and linear (blue) models. The extrapolated end times of constriction from these models ($t_{end}$) are compared to the departure time of PBP2B ($t_{2B}$). $t_{2B}$ is defined as the time point before the mNeonGreen-PBP2B signal drops to 50% of its maximum. (b) Difference between $t_{end}$ and $t_{2B}$ for both models for all ME7 cells. White circles, median; thick black lines, interquartile range; thin black lines, 1.5x interquartile range.

### SUPPLEMENTARY TABLES

**Supplementary Table 1:** Doubling times for various strains under different growth conditions. Errors are standard deviation.

| | **CDM 30°C** | **PHMM 30°C** | **CDM 37°C** | **PHMM 37°C** |
|---|---|---|---|---|
| **PY79** | 99.757 ± 10.730 | 49.171 ± 4.144 | 95.668 ± 9.897 | 35.872 ± 2.126 |
| **SH211** | 93.156 ± 6.602 | 59.659 ± 2.591 | 80.264 ± 4.592 | 42.304 ± 1.602 |
| **ME7** | 116.651 ± 26.788 | 58.938 ± 12.947 | 88.214 ± 10.577 | 35.573 ± 2.142 |
| **SH130** | 116.447 ± 23.174 | 48.488 ± 4.106 | 97.509 ± 10.451 | 39.184 ± 2.353 |
| **bGS28** [a] | 162.390 ± 16.818 | 49.999 ± 7.879 | 100.611 ± 18.670 | 37.876 ± 3.442 |
| **SH212** [a] | 151.631 ± 38.993 | 47.227 ± 17.515 | 119.269 ± 35.779 | 42.645 ± 2.832 |
| **SH212** [b] | 151.955 ± 16.587 | 50.832 ± 15.715 | 96.761 ± 23.822 | 43.653 ± 1.947 |
| **SH131** | 124.580 ± 3.448 | 62.934 ± 9.384 | 59.180 ± 3.035 | 45.141 ± 1.526 |
| **SH131** [c] | 147.992 ± 10.462 | 54.326 ± 3.843 | 76.646 ± 4.693 | 47.739 ± 1.804 |
| **bWM4** | 82.3046 ± 3.785 | 51.800 ± 2.367 | 94.225 ± 13.471 | 36.075 ± 1.405 |
| **bWM4** [d] | 81.0868 ± 9.881 | 52.259 ± 0.995 | 88.173 ± 10.648 | 35.863 ± 0.765 |

[a] 20 µMIPTG; [b] 20 µM IPTG, 0.08% xylose; [c] 10 µM IPTG; [d] 25 µM IPTG

**Supplementary Table 2**: Strain genotypes. All strains listed are *Bacillus subtilis* species.

| **Strain** | **Genotype/Description** [a] | **Reference/ Source** |
|---|---|---|
| PY79 | Prototroph | [1] |
| PB5250 | BS168 *trpC2 Δhag::kan* | [2] Bacillus Genetic Stock Centre |
| PL642 | JH642 *trpC2 pheA1 ftsZ::ftsZ-cam* | [3] |
| bAB215 | PY79 *amyE::erm*-$P_{hyperspank}$-*ftsAZ*(D213A) | [4] |
| bAB217 | PY79 *amyE::erm*-$P_{hyperspank}$-*ftsA-mNeonGreen*(SW)-*ftsZ*(D213A) | [4] |
| bGS28 | PY79 *pbp2B::erm*-$P_{hyperspank}$-*HaloTag-15aa-pbp2B* | [4] |
| bWM4 | PY79 *amyE::erm*-$P_{hyperspank}$-*ftsA-mNeonGreen-15aa-ftsZ* | [4] |
| ME7 | PY79 *pbp2B::mNeonGreen-15aa-pbp2B* | [4] |
| 2020 | 168 *trpC2* Ω *amyE::spc* $P_{xyl}$ *gfp-ftsZ* | [5] |
| SH130 | PY79 *Δhag ftsZ::ftsZ-gfp-cam* | This study |
| SH131 | PY79 *Δhag ftsZ::ftsZ-gfp-cam* Ω *amyE::erm*-$P_{hyperspank}$-*ftsAZ*(D213A) | This study |
| SH132 | PY79 *pbp2B::mNeonGreen-15aa-pbp2B* Ω *amyE::erm*-$P_{hyperspank}$-*ftsAZ*(D213A) | This study |
| SH211 | PY79 *Δhag* | This study |
| SH212 | PY79 *pbp2B::erm*-$P_{hyperspank}$-*HaloTag-15aa-pbp2B* Ω *amyE::spc* $P_{xyl}$ *gfp-ftsZ* | This study |

[a]Drug resistance cassettes: erm, erythromycin resistance; cam, chloramphenicol resistance; spc, spectinomycin resistance; kan, kanamycin resistance

**Supplementary Table 3**: Table of sample sizes. N.D indicates number of cells not determined; in conditions where B. subtilis forms multi-cellular filamentous chains it is not possible to determine the total number of cells used for a measurement without additional cellular labels.

| **Figure number** | **No. data points** | **No. cells** |
|---|---|---|
| 1a | 61 cells | 61 |
| 1b | 3256 data points | 61 |
| 1c nascent rings | 526 data points | 61 |
| 1c mature rings | 1053 data points | 61 |
| 1c constricting rings | 1677 data points | 61 |
| 1e-f | N.D | 766 |
| 1h nascent rings | 1066 filaments | 234 |
| 1h mature/ early constricting rings | 1588 filaments | 254 |
| 1h late constricting rings | 1538 filaments | 278 |
| 2c untreated | 438 filaments | 95 |
| 2c DMSO | 103 filaments | 23 |
| 2c PenG | 71 filaments | 11 |
| 2c PC19 | 180 filaments | 61 |
| 3b | 227 rings | 227 |
| 3d | 193 mature rings | 193 |
| 4b rich media 37C, untreated | 765 septa | 765 |
| 4b rich media 37C, PC19 | 419 septa | 419 |
| 4b poor media 30C, untreated | 373 septa | 373 |
| 4b poor media 30C, PC19 | 100 septa | 100 |
| 4c rich media 37C | 1178 septa | 1178 |
| 4c rich media 30C | 1018 septa | 1018 |
| 4c poor media 37C | 451 septa | 451 |
| 4c poor media 30C | 534 septa | 534 |
| 4d 37C Rich Media | 87 filaments | N.D. |
| 4d 30C Rich Media | 95 filaments | N.D. |
| 4d 37C Poor Media | 69 filaments | N.D. |
| 4d 30C Poor Media | 94 filaments | N.D. |
| Supplementary figure 2 PY79 30C poor media | 98 cells | 98 |
| Supplementary figure 2 SH130 30C poor media | 98 cells | 98 |
| Supplementary figure 2 PY79 37C poor media | 98 cells | 98 |
| Supplementary figure 2 SH130 37C poor media | 98 cells | 98 |
| Supplementary figure 2 PY79 37C rich media | 98 cells | 98 |
| Supplementary figure 2 SH130 37C rich media | 98 cells | 98 |
| Supplementary figure 4 | 61 cells | 61 |
| Supplementary figure 8a | 3256 data points | 61 |
| Supplementary figure 8b | 766 rings | 766 |
| Supplementary figure 10, ground truth (nascent FtsZ-GFP cells) | 1066 filaments | 234 |
| Supplementary figure 10, simulation ρ=1.3 filaments/septum | 603 filaments | 100 |
| Supplementary figure 10, simulation ρ=3.3 filaments/septum | 1446 filaments | 100 |

| | | |
|---|---|---|
| Supplementary figure 10, simulation ρ=6.5 filaments/septum | 2712 filaments | 100 |
| Supplementary figure 10, simulation ρ=37.5 filaments/septum | 2252 filaments | 80 |
| Supplementary figure 16, Wild-type | 87 filaments | N.D. |
| Supplementary figure 16, D213A | 37 filaments | N.D. |
| Supplementary figure 17d, Wild-type | 1635 data points | 60 |
| Supplementary figure 17d, D213A | 6009 data points | 229 |
| Supplementary figure 20, WT untreated | 765 septa | 765 |
| Supplementary figure 20, D213A | 256 septa | 256 |
| Supplementary figure 20, PC19 | 419 septa | 419 |
| Supplementary figure 24a, untreated | 493 septa | 493 |
| Supplementary figure 24a, DMSO | 188 septa | 188 |
| Supplementary figure 24a, PC19 | 110 septa | 110 |
| Supplementary figure 24b, untreated | 766 septa | 766 |
| Supplementary figure 24b, PC19 | 419 septa | 419 |
| Supplementary figure 27b, parabolic | 766 septa | 766 |
| Supplementary figure 27b, parabolic | 766 septa | 766 |

**Supplementary Table 4: Microscopy acquisition parameters.**

| **Figure number** | **Microscope configuration** | **Wavelength & power density** | **Exposure time & acquisition interval** | **Image pixel size** |
|---|---|---|---|---|
| **1a** | Custom inverted microscope, ring HiLO illumination | 488 nm 1-8 W/cm$^2$ | 1 second exposure, 1 frame/min | 65 nm |
| **1e** | Custom inverted microscope, ring HiLO illumination | 488 nm 1-8 W/cm$^2$ | 1 second exposure, no delay | 65 nm |
| **2b** | Custom inverted microscope, ring HiLO illumination | 488 nm 1-8 W/cm$^2$ | 1 second exposure, no delay | 65 nm |
| **3a** | Custom inverted microscope, ring HiLO illumination | 488 nm, 1-2 W/cm$^2$ | 1 second exposure, 1 frame/min | 65 nm |
| **3c** | Custom inverted microscope, ring HiLO illumination | 488 nm, 1-2 W/cm$^2$; 561 nm, 1-2 W/cm$^2$ | 1 second exposure, 1 frame/min | 65 nm |
| **4a-b** | Custom inverted microscope, ring HiLO illumination | 488 nm, 1-2 W/cm$^2$ | 1 second exposure, 1 frame/min | 65 nm |
| **4c** | Nikon NSTORM inverted microscope, HiLO illumination | 488 nm, 1.1 W/ cm$^2$ | 1 second exposure, 1 frame/ min | 64 nm |
| **4d** | Nikon NSTORM microscope, TIRF illumination | 488 nm 9.45 W/cm$^2$ | 1 second exposure, no delay | 64 nm |

| | | | | |
|---|---|---|---|---|
| **Supplementary figure 14** | Custom inverted microscope, ring HiLO illumination | 488 nm, 1-2 $W/cm^2$ | 1 second exposure, 1 frame/min | 65 nm |
| **Supplementary figure 15** | Custom inverted microscope, ring HiLO illumination | 488 nm, 1-2 $W/cm^2$ | 1 second exposure, 1 frame/min | 65 nm |
| **Supplementary figure 16** | Nikon NSTORM microscope, TIRF illumination | 488 nm 9.45 $W/cm^2$ | 1 second exposure, no delay | 64 nm |
| **Supplementary figure 17** | Custom inverted microscope, ring HiLO illumination | 488 nm, 1-2 $W/cm^2$ | 1 second exposure, 1 frame/min | 65 nm |
| **Supplementary figure 18** | Custom inverted microscope, ring HiLO illumination | 488 nm, 1-2 $W/cm^2$; 561 nm, 1-2 $W/cm^2$ | 1 second exposure, 1 frame/min | 65 nm |
| **Supplementary figure 19** | Custom inverted microscope, ring HiLO illumination | 488 nm, 1-2 $W/cm^2$ | 1 second exposure, 1 frame/min | 65 nm |
| **Supplementary figure 20** | Custom inverted microscope, ring HiLO illumination | 488 nm, 1-2 $W/cm^2$ | 1 second exposure, 1 frame/min | 65 nm |
| **Supplementary figure 24** | Custom inverted microscope, ring HiLO illumination | 488 nm, 1-2 $W/cm^2$ | 1 second exposure, 1 frame/min | 65 nm |

### SUPPLEMENTARY VIDEO LEGENDS

**Supplementary Video 1: FtsZ-GFP (SH130) organisation over cell cycle**. Representative video of FtsZ-GFP (SH130) dynamics, slow growth conditions (CDM, 30°C), imaged by ring-HILO microscopy with 1s exposure at 1 frame/ min. Video displayed at 20 frames per second (1200x actual speed). Scale bar, 1 µm.

**Supplementary Video 2: VerCINI imaging allows sensitive high-resolution imaging of division protein dynamics around the entire division septum for large numbers of cells.** Representative video of FtsZ-GFP (SH130) dynamics, imaged by VerCINI with ring-HILO illumination, 1s exposure and 2 second intervals. Unzoomed video section shows dynamics for all Z-rings recorded in a single VerCINI measurement. Zoomed video section highlights dynamics of 9 Z-rings in same measurement at higher magnification. Acquisitions displayed at 30 frames per second (60x actual speed).

**Supplementary Video 3: Exemplar results of VerCINI image processing on a nascent Z-ring for FtsZ-GFP (SH130) strain.** Representative video of FtsZ-GFP (SH130) dynamics, imaged continuously by VerCINI with ring-HILO illumination, 1s exposure. Left: Raw movie. Middle: Denoised and registered movie. Right: Movie after background subtraction using explicit model (Methods). Scale bar, 1 µm.

**Supplementary Video 4: FtsZ filaments in nascent Z-rings are sparse, highly dynamic and often static.** Representative videos of nascent FtsZ-GFP rings (SH130) imaged using VerCINI. Cells were imaged by ring-HiLO continuously with 1 s exposure. Acquisitions displayed at 15 frames per second(15x actual speed).Scale bar:1 µm.

**Supplementary Video 5: FtsZ filaments are predominantly treadmilling in mature/ early constricting rings**. Representative videos of mature/ early constricting FtsZ-GFP rings (SH130) imaged using VerCINI. Cells were imaged by ring-HiLO continuously with 1 s exposure. Acquisitions displayed at 15 frames per second (15x actual speed).Scale bar:1 µm.

**Supplementary Video 6: FtsZ filaments dynamics in late constricting rings are indistinguishable from those in mature/ early constricting rings.** Representative videos of mature/ early constricting FtsZ-GFP rings (SH130) imaged using VerCINI. Cells were imaged by ring-HiLO continuously with 1 s exposure. Acquisitions displayed at 15 frames per second (15x actual speed). Scale bar:1 µm.

**Supplementary Video 7: PC19 arrests FtsZ treadmilling within seconds throughout division.** Representative videos of FtsZ-GFP (SH130) cells imaged using microfluidic VerCINI during instantaneous perturbation with excess PC19. Cells were imaged by ring-HiLO continuously with 1 s exposure. Acquisitions displayed at 15 frames per second (15x actual speed). Scale bar: 500 nm.

**Supplementary Video 8: PenG does not affect FtsZ treadmilling dynamics.** Representative videos of FtsZ-GFP (SH130) cells imaged using microfluidic VerCINI during instantaneous perturbation with excess PenG. Cells were imaged by ring-HiLO continuously with 1 s exposure. Acquisitions displayed at 15 frames per second (15x actual speed). Scale bar: 500 nm.

**Supplementary Video 9: DMSO does not affect FtsZ treadmilling dynamics.** Representative videos of FtsZ-GFP (SH130) cells imaged using microfluidic VerCINI during instantaneous perturbation with 1% DMSO. Cells were imaged by ring-HiLO continuously with 1 s exposure. Acquisitions displayed at 15 frames per second (15x actual speed). Scale bar: 500 nm.

**Supplementary Video 10: PC19 treatment prevents nascent Z-rings from condensing or constricting.** FtsZ-GFP (SH130) cells imaged in a cellASIC device before and during treatment with excess PC19 in fast growth conditions (rich media 37$^{o}$C). Representative nascent ring (red circle) tracked manually using TrackMate. Cells were imaged by ring-HiLO with 1 s exposure at 1 frame/min. Acquisitions displayed at 10 frames per second (600x actual speed).

**Supplementary Video 11: PC19 treatment prevents some mature Z-rings from constricting.** FtsZ-GFP (SH130) cells imaged in a cellASIC device before and during treatment with excess PC19 in fast growth conditions (rich media 37°C). Representative mature ring (red circle) tracked manually using TrackMate. Cells were imaged by ring-HiLO with 1 s exposure at 1 frame/min. Acquisitions displayed at 10 frames per second (600x actual speed).

**Supplementary Video 12: Some mature Z-rings constrict post-PC19 treatment.** FtsZ-GFP (SH130) cells imaged in a cellASIC device before and during treatment with excess PC19 in fast growth conditions (rich media 37°C). Representative mature ring (blue circle) tracked manually using TrackMate. Cells were imaged by ring-HiLO with 1 s exposure at 1 frame/min. Acquisitions displayed at 10 frames per second (600x actual speed).

**Supplementary Video 13: Constricting Z-rings typically continue constricting post-PC19 treatment.** FtsZ-GFP (SH130) cells imaged in a cellASIC device before and during treatment with excess PC19 in fast growth conditions (rich media 37°C). Representative constricting ring (blue circle) tracked manually using TrackMate. Cells were imaged by ring-HiLO with 1 s exposure at 1 frame/min. Acquisitions displayed at 10 frames per second (600x actual speed).

**Supplementary Video 14: PenG rapidly stops constriction.** mNeonGreen-PBP2B (ME7) cells imaged in a cellASIC device before and during treatment with excess PenG in fast growth conditions (rich media 37°C). Cells were imaged by ring-HiLO with 1 s exposure at 1 frame/min. Acquisitions displayed at 10 frames per second (600x actual speed).

**Supplementary Video 15: Effect of PC19 treatment on FtsZ-GFP cells in slow growth conditions.** FtsZ-GFP (SH130) cells imaged in a cellASIC device before and during treatment with excess PC19 in slow growth conditions (poor media 30°C). Cells were imaged by ring-HiLO with 1 s exposure at 1 frame/min. Acquisitions displayed at 10 frames per second (600x actual speed).

**Supplementary Video 16: Effect of PC58 treatment on FtsZ-GFP cells.** FtsZ-GFP (SH130) cells imaged in a cellASIC device before and after treatment with excess PC58 in fast growth conditions (rich media 37°C). Cells were imaged by ring-HiLO with 1 s exposure at 1 frame/min. Acquisitions displayed at 10 frames per second (600x actual speed).

**Supplementary Video 17: Effect of GTPase-deficient mutant FtsZ(D213A) expression on Z-ring condensation and constriction.** A strain expressing FtsZ-GFP from the native locus and the GTPase-deficient mutant FtsZ(D213A) from an inducible promoter (SH131) imaged on agarose pads containing 10 µM IPTG in slow growth conditions (poor media 30°C). Liquid cultures were induced with 10 µM IPTG for 1 hour prior to imaging. Cells were imaged by ring-HiLO with 1 s exposure at 1 frame/min. Acquisitions displayed at 10 frames per second (600x actual speed).

**Supplementary Video 18: PC19 treatment prevents mature rings with low PBP2B signal from constricting.** GFP-FtsZ HaloTag-PBP2B (SH212) cells labelled with JF549 HaloTag ligand imaged in a cellASIC device before and during treatment with excess PC19 in fast growth conditions (rich media 37°C). Representative mature ring with low PB2B signal (white circle) tracked manually using TrackMate. Cells were imaged by ring-HiLO with 1 s exposure at 1 frame/min. Acquisitions displayed at 10 frames per second (600x actual speed). GFP-FtsZ: green, JF549-HaloTag-PBP2B: magenta.

**Supplementary Video 19: Mature Z-rings with high PBP2B signal continue constricting post-PC19 treatment.** GFP-FtsZ HaloTag-PBP2B (SH212) cells labelled with JF549 HaloTag ligand imaged in a cellASIC device before and during treatment with excess PC19 in fast growth conditions (rich media 37°C). Representative mature ring with high PB2B signal (white circle) tracked manually using TrackMate. Cells were imaged by ring-HiLO with 1 s exposure at 1 frame/min. Acquisitions displayed at 10 frames per second (600x actual speed). GFP-FtsZ: green, JF549-HaloTag-PBP2B: magenta.

**Supplementary Video 20: Effect of PC19 treatment on mNeonGreen-PBP2B cells in fast growth conditions.** mNeonGreen-PBP2B (ME7) cells imaged in a cellASIC device before and during treatment with excess PC19 in fast growth conditions (rich media 37°C). Cells were imaged by ring-HiLO with 1 s exposure at 1 frame/min. Acquisitions displayed at 10 frames per second (600x actual speed).

**Supplementary Video 21: Effect of PC19 treatment on mNeonGreen-PBP2B cells in slow growth conditions.** mNeonGreen-PBP2B (ME7) cells imaged in a cellASIC device before and during treatment with excess PC19 in slow growth conditions (poor media 30°C). Cells were imaged by ring-HiLO with 1 s exposure at 1 frame/min. Acquisitions displayed at 10 frames per second (600x actual speed).

### SUPPLEMENTARY NOTES

#### Supplementary Note 1: Tilted circle model provides estimates of Z ring diameters

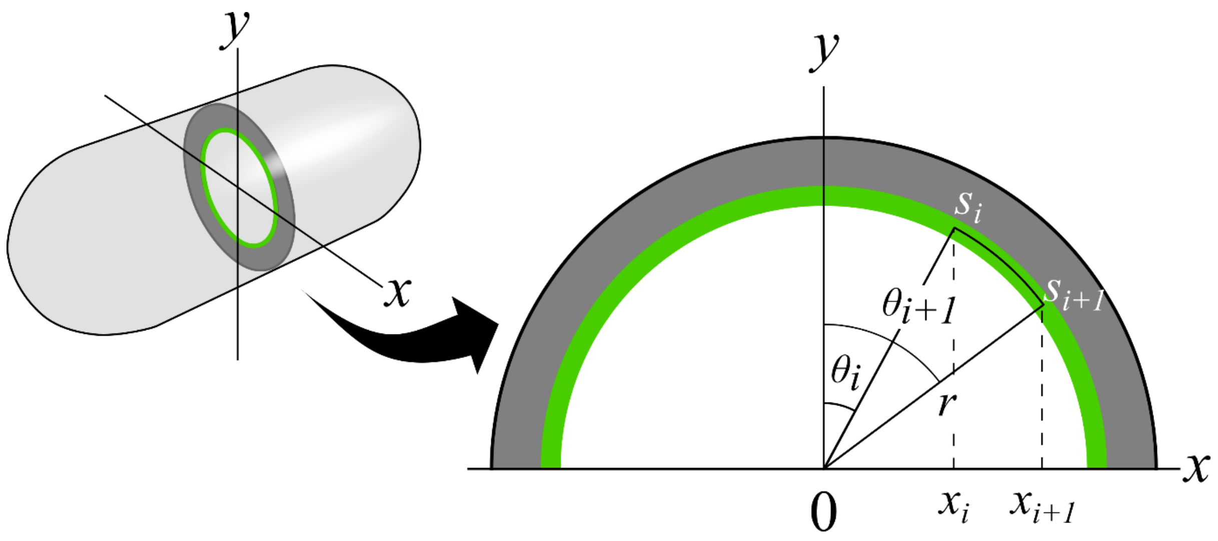


Consider a luminous circle of radius $r$ and uniform intensity that is rotated 90° about an axis going through its diameter. We want the intensity profile of the circle projected onto this axis. For simplicity, we first consider the top half of the circle. The fluorescence intensity $I(x)$ recorded between two points $x_i$ and $x_{i+1}$ will be proportional to the amount of luminous material in the circle directly above it, which is the arc length segment $\Delta s$:

$$I(x_i) = I_0(s_{i+1} - s_i)$$

Where $I_0$ is the intensity per unit length of the circle.

Defining the angle $\theta$ from the $y$-axis, $s_i = r\theta_i$, $x_i = r\sin\theta_i$, and so

$$s_i = r\sin^{-1}\left(\frac{x_i}{r}\right)$$

So the arc length segment can be described in terms of $x$ as

$$s_{i+1} - s_i = r\left[\sin^{-1}\left(\frac{x_{i+1}}{r}\right) - \sin^{-1}\left(\frac{x_i}{r}\right)\right]$$

And so the intensity of the full circle in slice $i$ is

$$I(x_i) = 2I_0 r\left[\sin^{-1}\left(\frac{x_{i+1}}{r}\right) - \sin^{-1}\left(\frac{x_i}{r}\right)\right]$$

Since the depth of field in high-NA microscope images is very small, the intensities of real images will be lower near the centres of their line profiles than the model predicts. As a result, this model can be expected to produce a slight overestimate of the septal diameter in such images.

**Supplementary Note 2: Constant septal PG synthesis rate predicts parabolic septal constriction**

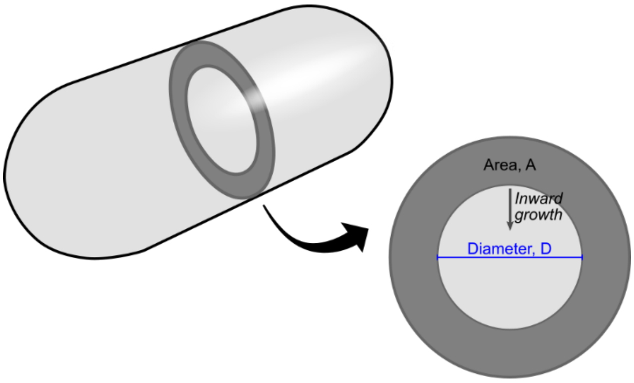


Consider a model where new cell wall material is added to a septal disk of surface area *A,* unconstricted cell diameter $D_0$, diameter of septal leading edge *D,* and fixed thickness at constant septal synthesis rate *k* (units [Area]/[Time]):

$$\frac{dA}{dt} = k.$$

Since $A = \frac{\pi}{4}(D_0^2 - D^2)$, $\frac{dA}{dD} = -\frac{\pi}{2}D$, and by the chain rule:

$$\frac{dD}{dt} = -\frac{2k}{\pi D}.$$

Integrating with respect to time gives:

$$D(t) = \sqrt{D_0^2 - \frac{4k}{\pi}t}.$$

This constant PG synthesis model predicts that constriction will accelerate parabolically with time as a result of septal geometry, consistent with the data. An area $kdt$ deposited around a large septum will correspond to a small decrease in diameter, while the same area added to a small septum corresponds to a larger decrease in diameter.

*Linear constriction model.* Alternatively, a constant constriction rate model can be defined:

$$D(t) = D_0 - at,$$

where *a* is a rate constant. By comparison with previous analysis, a constant diameter constriction rate would imply that septal synthesis around the septum leading edge slows down over time, as the diameter of the septum decreases.

Comparison of the two models indicates that the parabolic model is most consistent with the data (Supplementary Figure 19).

**Supplementary Note 3: Classification of Z-ring phase for horizontal and vertical cell microscopy data**

Z-ring organisation was measured by conventional time lapse microscopy of horizontally immobilised cells (Figure 1a). Upon initial assembly, FtsZ filaments form a diffuse *nascent Z-ring* structure at midcell. After gradually increasing in intensity, the Z-ring rapidly condenses into a dense, narrow band, which we term a *mature Z-ring,* followed after some time by constriction initiation (*constricting Z-ring)*. Z-ring diameter and thickness were measured across division for each septum (Figure 1b). A parabolic constriction model was fitted to the Z-ring diameter time trace (Methods), and all trajectories aligned such that fitted time of constriction initiation is *t=0*.

Septal density was calculated for all time points, defined as the total septal intensity divided by the fitted Z-ring circumference. Septal density was calculated instead of the simpler total intensity quantity as this quantity was directly comparable to septal intensities measured using VerCINI.

Inspection of the ring thickness plots (Figure 4b) and a plot of ring thickness versus diameter (Figure 4c) indicated that nascent Z-rings initially are initially decondensed, with large, variable axial thickness, consistent with their composition of sparse FtsZ filaments. Nascent Z-rings then rapidly condensed to form dense Z-rings (Figure 4c), close to the thickness of a diffraction limited structure (approximately 300 nm for this system and sample). This qualitative analysis was supported by automated step-detection in the axial thickness time series which was used to classify axial thickness into either one or two states (Methods). For time traces where two states were detected, we found that the earlier detected state was high variance and larger axial thickness (Figure 1b, Figure 21), consistent with the decondensed state label. The later detected state had low variance with observed thickness 300-350 nm, consistent with the condensed state label. In axial thickness time series where only one state was detected, the axial thickness observed was < 400 nm, and qualitatively could not be distinguished from the condensed Z-ring state identified in two-state time traces. We concluded that these single state trajectories represent Z-rings which had also condensed by the time of detection. Transitions between the decondensed and condensed states were observed to occur stochastically, with a broad spread of condensation times, median condensation time 11 minutes [IQR 8-17] before constriction initiation (Fig 21b). State detection supports the conclusion that Z-rings initially assemble as sparse decondensed rings before rapidly condensing into a narrow band for substantial time prior to constriction initiation.

Z-rings were classified as “Nascent”, “Mature” and “Constricting” as follows (Figure 1b-c):

- *Nascent Z-rings*: ring thickness detected as decondensed via step-finding, not constricting *(time after initiation < 0)*
- *Mature Z-rings*: ring thickness detected as condensed via step-finding, not constricting *(time after initiation < 0)*
- *Constricting Z-ring*: actively constricting *(time after initiation > 0)*

Classification of VerCINI Z-ring phase is harder than horizontal time lapse data because it lacks the long term time-lapse and high-resolution axial thickness information required to trivially assign Z-ring state. We observed that septal density steadily increases throughout cell division (Figure 1b). We thus investigated if septal density could be used, together with Z-ring diameter, to determine Z-ring phase for VerCINI data based on a calibration from the horizontal cell data.

We plotted septal density against diameter for all horizontal cell data points classified into Z-ring phase using the previous analysis of axial thickness and constriction state (Supplementary Figure 6a). We observed that significantly constricted Z-rings are trivially separable from other Z-ring phases based on diameter alone (Supplementary Figure 6a; constricting Z-rings normalized diameter < 0.9, purple circles). Unconstricted mature and nascent Z-rings (Supplementary Figure 6a; cyan circles, mature Z-rings; blue circles, nascent Z-rings), together with constricting Z-rings that had not yet constricted a large amount (Supplementary Figure 6a; constricting Z-rings normalized diameter > 0.9, purple circles), showed a large distribution of fitted diameters around 1 normalized diameter.

We made a kernel density plot of septal density for all Z-rings with observed normalized diameter > 0.9 in order to determine whether phase of unconstricted Z-rings could be separated by intensity alone (Supplementary Figure 6a, right). We observed that the intensity distribution for each Z-ring phase formed distinct peaks, albeit with significant overlap based on horizontal cell measurements only.

We next examined whether the increased intensity resolution gained by VerCINI would be sufficient to separate Z-rings into their distinct stages (Supplementary Figure 6b). We plotted median septal intensity at the septum for each Z-ring against fitted Z-ring diameter for cells imaged by VerCINI and observed a similar intensity-diameter distribution to the horizontal cell data.

As for the horizontal cell analysis, significantly constricted Z-rings were separable from other data points based on diameter: these we classified as *late constricting* (diameter < 800 nm). We calculated kernel density of all Z-ring septal intensities with a diameter greater than 800 nm (Supplementary Figure 6b, right, red line). Consistent with the horizontal cell measurements we observed a broad distribution with 3 overlapping peaks. Based on the horizontal cell results we fitted 3 Gaussian peaks to the data, in order to estimate the degree of overlap of the nascent, mature and constricting Z-ring intensity distributions (Supplementary Figure 6b, purple, cyan, magenta dashed lines). The mature and nascent cell populations showed greater intensity separation in the VerCINI separation than the horizontal cell data, consistent with the presence of a large diameter, dim intensity cluster in the intensity-diameter plot of VerCINI data (Supplementary Figure 6b, left).

Placing a threshold at the 500 (arbitrary units) intensity crossover point of the mature and nascent populations allowed robust classification of different cell division phases in the VerCINI data with little overlap, as estimated from the 3 fitted populations: *nascent Z-rings* (84% nascent Z-rings, 16% mature Z-rings and 0% early constricting Z-rings) and *mature/early constricting Z-rings* (62% mature Z-rings, 27% early constricting Z-rings, 10% nascent Z-rings). Based on this analysis, we classified VerCINI Z-ring state as follows:

- *Nascent Z-rings*: ring diameter > 800nm, septal intensity < 500,
- *Mature/early constricting Z-rings*: ring thickness > 800nm, septal intensity > 500,
- *Constricting Z-ring*: ring thickness < 800nm, septal intensity > 500.

Supporting the conclusion that Z-rings can be classified in VerCINI based on diameter and septal intensity, observed spatiotemporal organisation of classified Z-rings was consistent with morphology from horizontal cell time-lapses. *Nascent Z-rings* were patchy and dynamic, often containing large gaps (Figure 1e, Supplementary Figure 5a). *Mature/early constricting* and *late constricting* Zrings were dense and continuous to the diffraction limited resolution of the measurement (Figure 5b-c).

**Supplementary Note 4: Investigation of robustness of measured FtsZ filament properties to changes in FtsZ filament density**

We performed simulations to investigate how FtsZ filament speed, lifetime, processivity and the observed fraction of immobile FtsZ filaments depended on FtsZ filament density in the Z-ring (Methods, Supplementary Figure 10). The principle of the simulations was to simulate Z-rings containing FtsZ filaments with dynamics identical to those observed in nascent Z-rings at low filament density, and to gradually increase the filament density over a large range to investigate the degree to which this introduced bias in the measured parameters, and specifically to investigate whether the filament dynamics measured in mature or constricting FtsZ rings were likely to be reliable.

FtsZ filament density of simulated nascent Z-rings was chosen to match the observed number of manually traced filaments in nascent Z-rings, as the filament density there was low enough that filament detection bias was expected to be low. To minimize undercounting bias, the density of FtsZ filaments in mature Z-rings was estimated by multiplying the estimated nascent Z-ring density by 2.6, the observed ratio of total Z-ring intensity for mature and nascent Z-rings (Supplementary Figure 8b). The accuracy of the FtsZ filament density estimate used for the mature and nascent Z-ring simulations assumes that both the manual estimate of FtsZ filament density in nascent Z-rings and the VerCINI Z-ring intensity measurements are accurate. Additionally, the filament density estimate in mature Z-rings also assumes that observed Z-ring intensity is proportional to the number of FtsZ filaments, which could become inaccurate if FtsZ filament length changes. To control for any errors in density estimation, additional simulations at 2x and 12x the estimated mature Z-ring filament densities were performed. This allowed us to investigate the extent to which large changes to FtsZ filament density, even beyond the expected FtsZ filament density, could perturb observed FtsZ filament speeds and other measured filament parameters. Simulated FtsZ ring time lapse movies were analysed identically to the experimental data, with FtsZ filament speed estimated by kymograph analysis.

Increased FtsZ filament density caused an approximately linear decrease in the fraction of detected filaments but had little effect on observed FtsZ filament speeds, except for a modest increase in observed FtsZ filament speed at the highest simulated FtsZ filament density. Notably, the fraction of detected immobile FtsZ filaments, i.e. the ability to robustly detect and differentiate immobile and treadmilling FtsZ filaments remained accurate even at the highest simulated FtsZ filament density (Supplementary Figure 10Ci,D). At high simulated FtsZ density, we found that FtsZ filament lifetime and processivity are underestimated.

Based on these simulations we conclude that FtsZ filament speeds and immobile filament fractions can be measured accurately across a wide range of FtsZ filament densities encompassing typical experimental conditions. However the experimental measurements of FtsZ filament lifetime and processivity in mature/ constricting Z-rings may somewhat underestimate the real values.
